# GC content across insect genomes: phylogenetic patterns, causes and consequences

**DOI:** 10.1101/2023.09.11.557135

**Authors:** Riccardo G. Kyriacou, Peter O. Mulhair, Peter W. H. Holland

## Abstract

The proportions of A:T and G:C nucleotide pairs are often unequal and can vary greatly between animal species and along chromosomes. The causes and consequences of this variation are incompletely understood. The recent release of high-quality genome sequences from the Darwin Tree of Life and other large-scale genome projects provides an opportunity for GC heterogeneity to be compared across a large number of insect species. Here we analyse GC content along chromosomes, and within protein-coding genes and codons, of 150 insect species from four holometabolous orders: Coleoptera, Diptera, Hymenoptera, and Lepidoptera. We find that protein-coding sequences have higher GC content than the genome average, and that Lepidoptera generally have higher GC content than the other three insect orders examined. GC content is higher in small chromosomes in most Lepidoptera species, but this pattern is less consistent in other orders. GC content also increases towards subtelomeric regions within protein-coding genes in Hymenoptera, Coleoptera and, most strikingly, Lepidoptera. Two species of Diptera, *Bombylius major* and *B. discolo*r, have very atypical genomes with ubiquitous increase in AT content, especially at third codon positions. Despite dramatic AT-biased codon usage, we find no evidence that this has driven divergent protein evolution. We argue that the GC landscape of Lepidoptera, Hymenoptera and Coleoptera genomes is influenced by GC-biased gene conversion, strongest in Lepidoptera, with some outlier taxa affected drastically by counteracting processes.

## Introduction

Animal genomes can display non-homogeneous distributions of DNA bases, where the local percentage of adenine (A) and thymine (T) content is not equal to that of guanine (G) and cytosine (C). Whilst some level of small-scale nucleotide heterogeneity is expected across a genome, particularly within structural and repetitive DNA sequences, larger GC stretches (approximately at the 100-kb scale) have repeatedly been observed within animal genomes (Figuet et al., 2014; Romiguier et al., 2010; Weber et al., 2014). Nucleotide heterogeneity stretches also include coding sequences (CDS) in the genome, suggesting that the primary processes underpinning GC variation are independent of sequence function. In the most extreme cases, documented in gerbils and in birds, GC-rich regions of the genome have affected the ability to accurately sequence DNA, leading to erroneous reports of missing genes (Benjamini and Speed, 2012; Hargreaves et al., 2017; Hron et al., 2015). In addition to variation along a genome or a chromosome, individual taxa have been noted with unusually high or low overall GC content; for example, the honeybee *Apis mellifera* (33% GC; Weinstock et al., 2006), hydroid *Hydra magnipapillata* (29% GC; Chapman et al., 2010) and sea lamprey *Petromyzon marinus* (46% GC, rising to 75% GC at third codon positions; Smith et al., 2013). The evolutionary basis and consequences of GC differences between species need further study through deep sampling of diverse taxa.

Nucleotide heterogeneity is shaped by an interplay between several factors including mutational bias (Boulikas, 1992; Kotari et al., 2023; Wolfe et al., 1989), base-specific excision repair of DNA mismatches (Krokan and Bjørås, 2013), selection (Eyre-Walker and Hurst, 2001), and genetic drift (Bulmer, 1991). An important force is thought to be the meiosis-associated process of GC-biased gene conversion (gBGC) (Duret and Galtier, 2009; Figuet et al., 2014; Marais, 2003; Pessia et al., 2012; Romiguier and Roux, 2017). If strand invasion at meiosis occurs over a region with a heterozygous site, the mismatch repair process can alter one of the bases to generate an A:T or G:C complement. This process has a biochemical bias in converting the mismatched site in the double helix towards G:C pairs (Duret and Galtier, 2009; Halldorsson et al., 2016). Hence, in regions of the genome with high frequency of recombination, a build-up of GC can occur over time, unless strongly selected against (Kostka et al., 2012). The process of gBGC is thought to underpin the existence of localised regions of high GC in gerbil and bird genomes (Brekke et al., 2023; Hargreaves et al., 2017; Hron et al., 2015; Pracana et al., 2020), and the observation that GC content is negatively correlated with chromosome length in several vertebrates (Goodstadt et al., 2007; Matsubara et al., 2012; Pessia et al., 2012). The latter is linked to the fact that at meiosis, chromosome pairs must undergo at least one crossover; hence DNA sequences on smaller chromosomes are affected by gBGC more frequently (Figuet et al., 2014; Romiguier et al., 2010; Saito and Colaiácovo, 2017). It is, however, unclear if the ‘small chromosome rule’ applies more widely across the animal kingdom, or indeed if many other animals besides gerbils and birds have local peaks of high GC driven by gBGC.

Investigations into GC heterogeneity have been plagued by inadequate taxonomic sampling (Romiguier et al., 2010); in insects this had been further compounded by a paucity of high quality insect genome assemblies (Li et al., 2019; but see Wright et al. 2023). Yet isolated studies on insect genomes yield intriguing results. Superimposed on the overall low GC percentage of the honeybee genome is a bimodal pattern of GC content (Jørgensen et al., 2007; Kent and Zayed, 2013), which like the gerbil and bird examples caused genes to be missed during genome sequencing (Elsik et al., 2014). In addition, at least in one lepidopteran genus (*Leptidea*), evidence for gBGC has recently been reported (Boman et al., 2021; Näsvall et al., 2023b). The recent advent of long-read DNA sequencing, coupled with Hi-C sequencing, has enabled the rapid generation of very high quality genome assemblies. The Darwin Tree of Life (DToL) is one of several projects exploiting these technologies to sequence and assemble high-quality genomes from large numbers of species (Crowley et al., 2023; The Darwin Tree of Life Project Consortium, 2022; Wright et al. 2023). Here we interrogate 150 newly generated insect genome assemblies to investigate GC sequence heterogeneity across Lepidoptera, Diptera, Hymenoptera, and Coleoptera. We ask whether there are consistent differences in GC content between orders, whether GC content is affected by chromosome size or chromosome position in insects, and whether patterns of GC heterogeneity extend to coding sequences, potentially influencing protein evolution.

## Results

### GC content of insect genomes is higher in coding regions

Using genome sequences for 150 insect species, we performed a genome-wide investigation of GC content, asking if GC and AT percentages differ between coding genes and the whole genome. We also examined 3^rd^ codon positions (GC3) as differences in GC3 can be used as an indicator of gBGC or mutational bias since this metric largely overcomes the influence of selection for amino acids encoded by GC-rich codons such as alanine, glycine, and proline. For GC3 analysis we included two important data filtering steps. First, we identified a set of single copy orthologous genes (SCO) found across the insect order of interest (772 SCOs for Lepidoptera, 849 for Diptera, 1330 for Coleoptera, and 1944 for Hymenoptera). Second, we aligned and trimmed the deduced SCO protein sequences to ensure that only homologous codons were compared (1,251,024 codons for Lepidoptera, 1,439,484 for Diptera, 3,430,818 for Hymenoptera and 2,103,432 for Coleoptera). These filtering steps were included to remove the influence of species-specific gene duplications, spurious protein coding gene predictions, artefactual frameshifts, and errors in exon/intron prediction, any of which could distort comparisons of GC3. The three metrics were calculated from the complete genomes of 60 Lepidoptera, 42 Diptera, 33 Hymenoptera and 15 Coleoptera (Figure 1; Supplementary Data).

**Figure 1.**
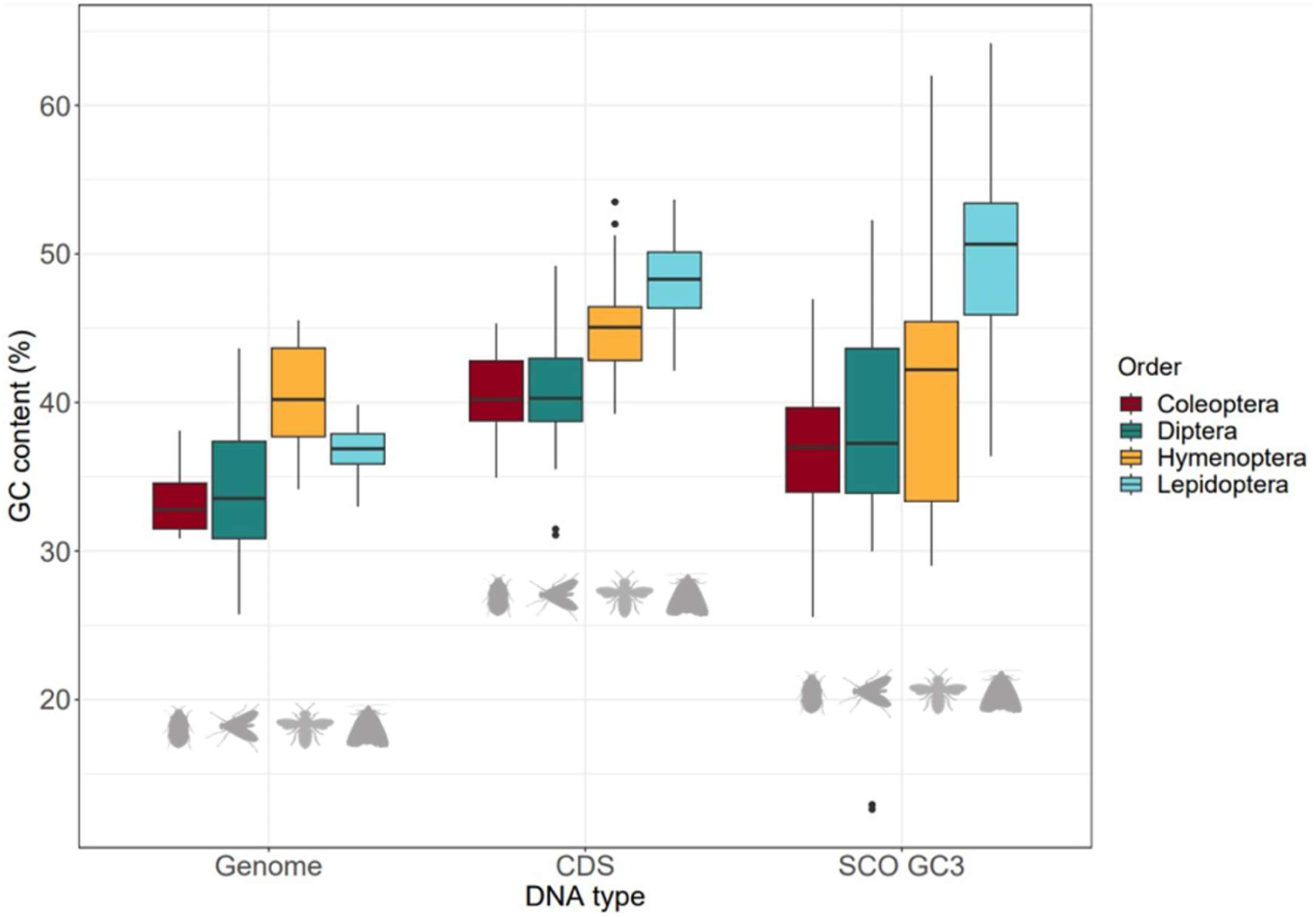
Percentage GC in complete genome, coding sequences (CDS) and third codon positions of single copy orthologs (SCO GC3). Data plotted for 60 Lepidoptera, 42 Diptera, 33 Hymenoptera and 15 Coleoptera species; each point plotted is a species. Box edge values correspond to first and third quartiles. Upper whisker extends to the largest value 1.5x interquartile range. Lower whisker extends to the smallest value 1.5x interquartile range. Data points beyond these limits are represented as outlier points.

A two-way analysis of variance (ANOVA) was performed using a linear model with DNA type (genome vs. CDS vs. GC3) and insect order as main factors. Significant effects on GC content were observed for DNA type (F_2, 438_ = 118.260, P < 2.2e-16) and order (F_3, 438_ = 65.713, P < 2.2e-16). The interaction between order and DNA type was also significant (F_6,_ 438 = 12.388, P < 6.6e-13). To identify the comparisons underlying these differences, a TukeyHSD post-hoc test was performed. This showed that for insect orders combined, GC content is significantly higher within CDS regions than across the entire genome (P =0, Supplementary table S1). The same is evident from examining third codon positions only, although this is from a smaller dataset due to filtering for SCOs. There is no significant difference between GC content of CDS regions and ortholog third codon positions (Supplementary Table S1). Hymenoptera have the highest average GC content across the entire genome (Figure 1), with the distribution significantly higher than in Diptera or Coleoptera (Supplementary table S1). In contrast, Lepidoptera have the highest average GC content within coding genes; the distribution for CDS regions is significantly greater than in Diptera or Coleoptera, and GC3 distribution significantly higher than in the other three orders (Supplementary Table S1). In summary, GC content is highest within coding regions of insect genomes, Hymenoptera have the highest genome-wide GC content, and Lepidoptera have the highest GC content within coding genes.

### Phylogenetic patterns of GC content reveals outlier lineages

To ascertain if patterns of GC evolution are consistent within an order, and to detect outlier lineages, we plotted GC against phylogenetic trees for each order (Figure 2; Supplementary Figure S1); trees were inferred using Orthofinder (Emms and Kelly, 2019). For each order, we plotted GC content for the genome (outer ring), coding regions (middle ring), and third codon positions of SCOs (inner ring, Figure 2).

**Figure 2.**
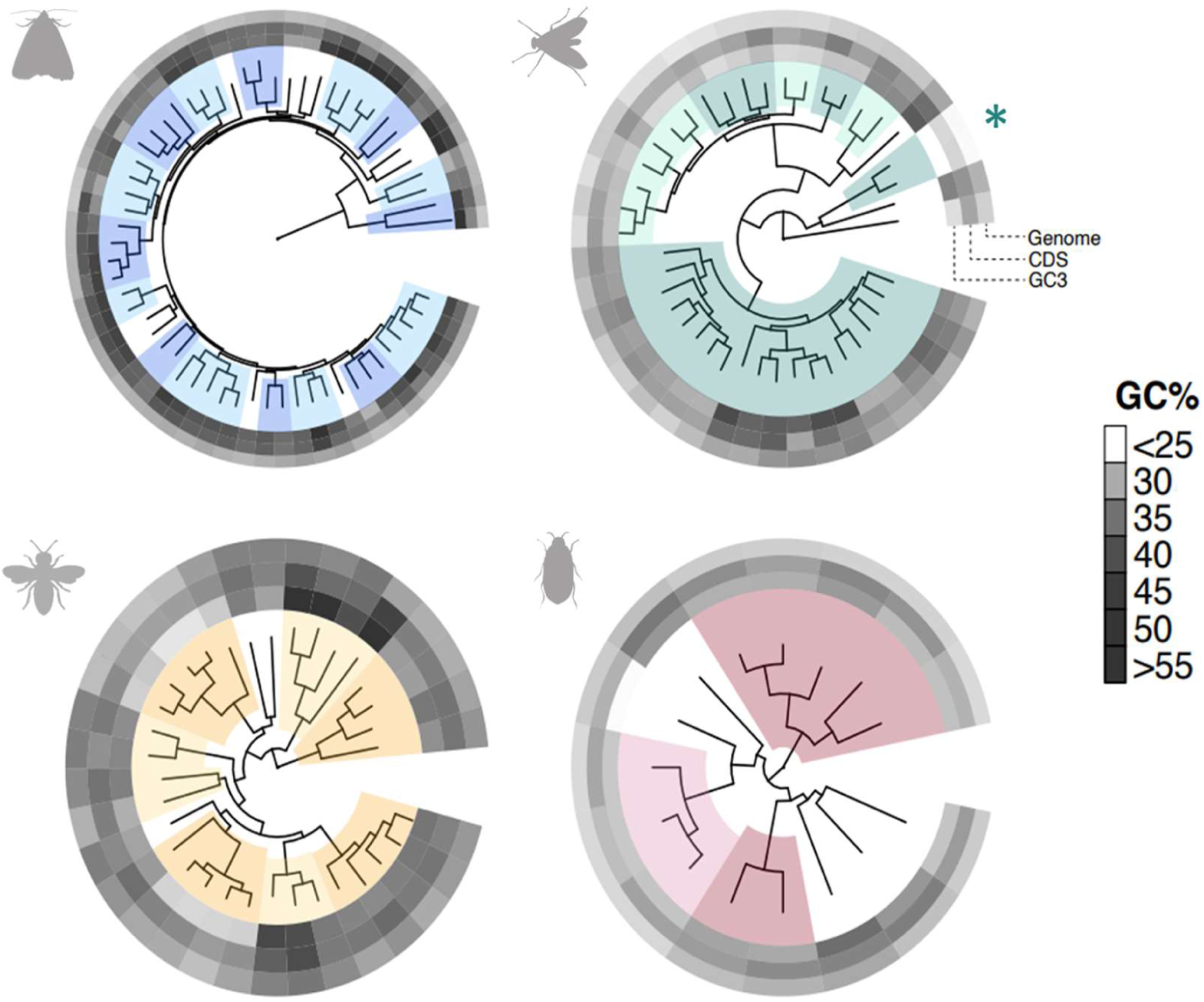
Phylogenetic plots comparing GC percentage across the genome (outer circle), in coding genes (middle circle) and at third codon position of single copy orthologs (SCO GC3; inner circle). For each order, families with two or more representative species are alternatively highlighted. Lepidoptera in blue, Diptera in green, Hymenoptera in yellow, Coleoptera in red. In the Diptera tree, Bombyliidae species are labelled with an asterisk.

The phylogenetic patterns show that the higher GC content in coding regions and at third codon positions, noted above for Lepidoptera, is seen across the phylogenetic diversity of this insect order. Indeed, most patterns of GC content are relatively consistent within each order. There are, however, some outlier species and genera. Hymenoptera see the greatest intra-order variation in GC content. We note higher GC3 in the family Ichneumonidae, a group of parasitic wasps consisting of *Ophion luteus* (59.5%), *Buathra laborator* (51.5%), *Ichneumon xanthorius* (59.8%), and *Amblyteles armatorius* (62.0%) in our dataset (Figure 2; Supplementary Figure S1). In contrast, we see lower SCO GC3 in the Vespidae family and the four bumblebee species in our data set. In the wasps this ranges from 29.0% in *Vespula vulgaris* to 33.3% in *Dolichovespula saxonica*, and in bumblebees this ranges from 31.2% in *Bombus terrestris* to 32.3% in *Bombus hortorum*. In Coleoptera we note aberrantly low GC3 (25.6%, relative to mean 36.7% across Coleoptera) in the click beetle species *Agrypnus murinus*, but a general consistency across the order. Finally, in Diptera we find highest GC content in the clade of Eristalini plus Milesiini (six species in our dataset, SCO GC3 ranging from 48.3-52.3%) within the hoverfly family (Syrphidae), as well as in the Small beegrabber fly *Thecophora atra* (49.6% GC3). The most striking outlier is the extremely low GC percentage, across all three DNA categories, for the two bee-fly species (family Bombyliidae); *Bombylius major* and *B. discolor* (green asterisk in Figure 2). Specifically, the Dark-edged bee-fly *B. major* has a genome GC content of 26.0%, coding sequence GC of 31.1%, and single copy ortholog GC3 of just 12.6%. The Dotted bee-fly *B. discolor* has similarly low values: genome GC 25.7%, CDS GC 31.5% and, SCO GC3 12.9%.

### GC content is related to chromosome size in some insects

In several vertebrate taxa, smaller chromosomes have a higher GC content, thought to be related to recombination frequency per site (Goodstadt et al., 2007; Matsubara et al., 2012). To test if a similar inverse relationship is also observed in insects, we plotted genome GC data against chromosome size for each order (Figure 3A). On these pooled data, negative regressions are not significant for Lepidoptera, Coleoptera, and Hymenoptera, but are significant for Diptera (slope = -9.666e-09, P < 2.2e-16; Supplementary Table S2). We note, however, that chromosome sizes vary greatly within an order, especially Diptera (Supplementary Figure S2), so combining species may mask species-specific trends. Plotting the data for species separately reveals that most Lepidoptera species (48/59) show a significant inverse correlation between chromosome size and GC content (Figure 3B; Supplementary Data). By contrast, relatively few species of Coleoptera, Diptera or Hymenoptera show such a trend: 5/15, 12/42, 4/32 species respectively (Figure 3B; Supplementary Data). These results imply that, in holometabolous insects, smaller chromosomes do not always have higher GC content, although the ‘small chromosome effect’ is most widespread in Lepidoptera.

**Figure 3.**
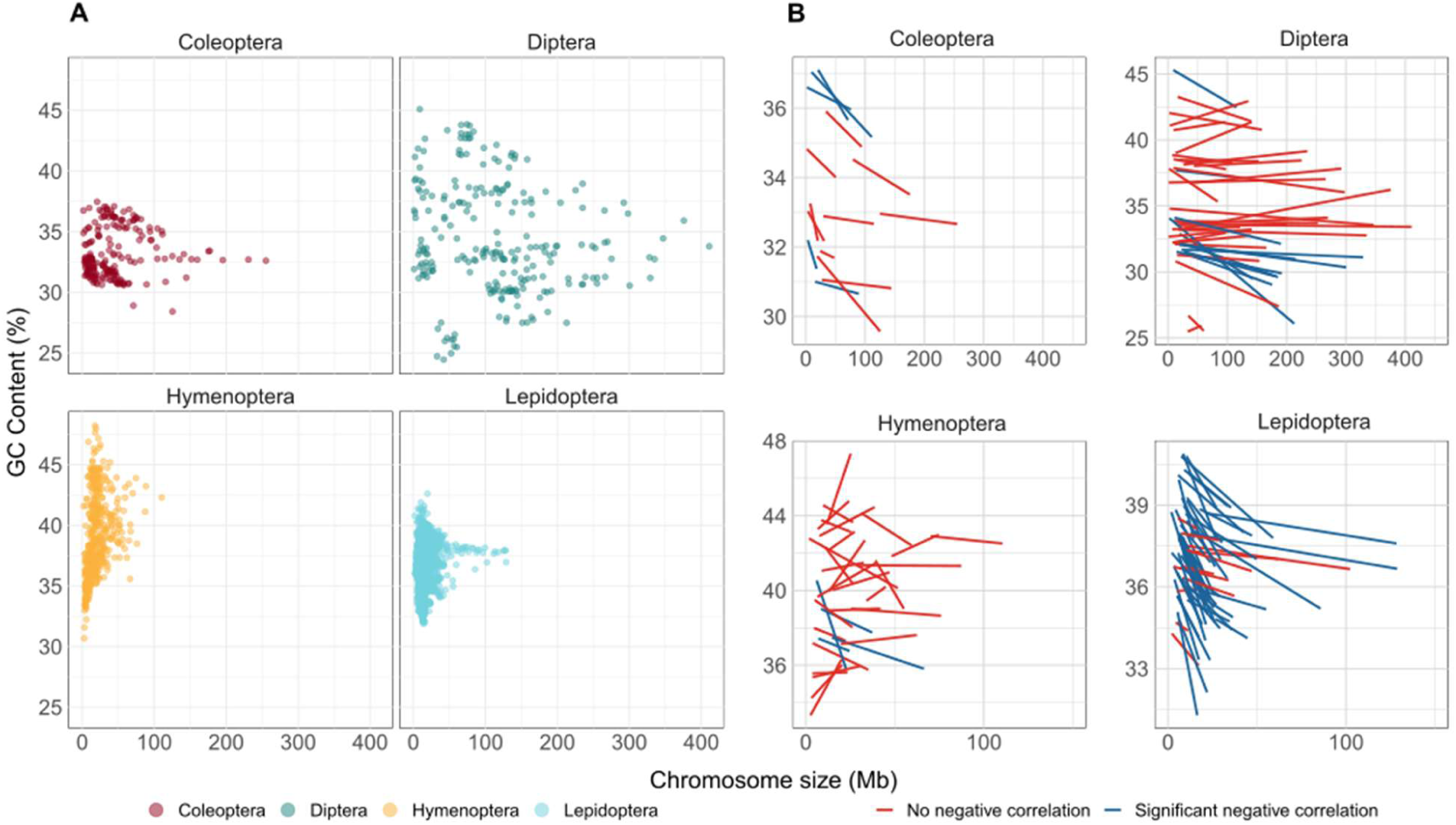
(A) GC content against chromosome size plotted for four insect orders. Each point represents GC content and size for one chromosome in one species. (B) GC content against chromosome size plotted for insect species separately. For each species, linear plots best fit are shown. Blue lines denote significant negative correlation (Pearson’s correlation coefficient, P < 0.05). Lines are red if negative correlations were not significant or for positive correlations. Unplaced scaffolds are omitted, as are two species where genome assemblies are not chromosome-level (*Bombus terrestris* and *Neomicropteryx cornuta)*.

### GC content decreases with distance from telomere in three insect orders

Recombination frequency is reported to increase towards the telomeric ends of chromosomes in various taxa (Coop and Przeworski, 2007; Haenel et al., 2018; Ma et al., 2015; Mouresan et al., 2019; Rockman et al., 2010). This is predicted to cause increased gBGC, and hence higher GC content towards telomeres (Arndt et al., 2005; Duret and Galtier, 2009). To test this effect in insects, we examined GC3 of SCO genes in relation to distance from telomere for each insect order separately (Figure 4A; Supplementary Table S3; Supplementary Data). We conducted the analysis on SCO GC3 to avoid confounding factors such as repetitive DNA, telomeric sequences, or repeated genes, and to ensure equivalent data are compared between species. We recognise that each orthologous gene may occupy a different chromosomal position in each species; hence we used mean distance from telomere and mean GC3 value for each gene.

**Figure 4.**
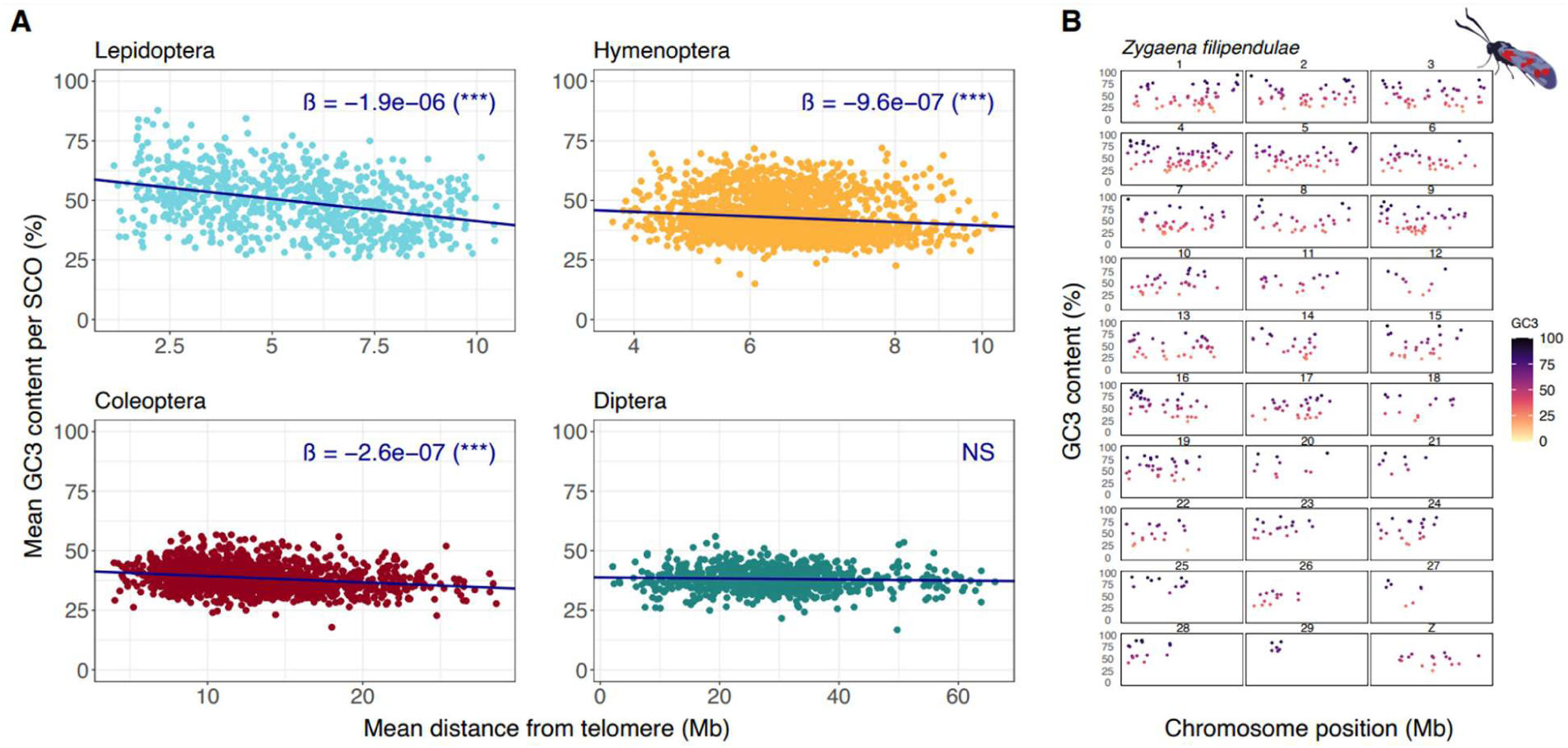
(A) GC3 content (%) against distance from telomere (Mb) plotted for 772 Lepidopteran, 1944 Hymenoptera, 1330 Coleoptera, and 849 Diptera single copy orthologs (SCOs). For each SCO (i.e. each point in the plot), mean GC3 and mean distance from the telomeres was calculated for each order containing 60, 33, 15, and 42 species respectively. Regression slopes superimposed for each order along with significance level (*** = P < 0.01, NS = Non-significant). (B) Chromosomal landscape plot showing change in GC3 content (y-axis) along chromosomes (x-axis) for single copy orthologs in a lepidopteran; Six-spot burnet moth *Zygaena filipendulae* is used as an example. Each panel represents a chromosome.

We find significant support for a negative correlation between GC3 and distance from telomeres within Lepidoptera, Hymenoptera, and Coleoptera (P < 6.32e-08; Supplementary Table S3). Hence, coding genes closer to telomeres, where recombination is expected to be increased, are higher in GC3 than those further away, consistent with biased gene conversion. Lepidoptera showed the steepest regression slope, a factor of 10 steeper than in Hymenoptera and Coleoptera. However, the ‘telomere-effect’ is not the main driver of the variance in GC3 content, as indicated by the low R squared values generated by the regressions (Lepidoptera R^2^: 0.0459, Hymenoptera R^2^: 0.01445, Coleoptera R^2^: 0.0459). Despite the telomere effect being a weak force overall, it has driven detectable patterns within some Lepidoptera chromosomes. Analysing GC3 against position for all SCOs in a single species (*Zygaena filipendulae*), chromosome GC3 landscapes can be constructed with noticeable increases of GC3 towards the ends of telomeres (Figure 4B). These results are consistent either with gBGC being a stronger force in Lepidoptera, or recombination being more often telomere-associated in Lepidoptera.

### AT-rich genomes of *Bombylius* have skewed codon usage but not unusual protein evolution

The strikingly high AT (low GC) content of bee-flies, family Bombyliidae, was noted above (asterisk in Figure 2). To test whether the high AT content extends to all genes, we plotted GC3 for all single copy orthologs for the 42 Dipteran whole genomes studied (Figure 5A). The majority of Dipteran SCOs have a relatively stable GC3 around 30-40% (Figure 5A); in *B. major* and *B. discolor* the orthologous genes have GC3 below 15%, averaging just 12.6 and 12.9%, respectively. Furthermore, plotting GC3 of SCOs along *Bombylius* chromosomes, shows that GC3 does not vary greatly with position along the chromosome (Figures 6B & C). These analyses reveal that the drive to very high AT (low GC) extends across protein coding genes, across chromosomes, and occurred on the lineage leading to Bombyliidae.

**Figure 5.**
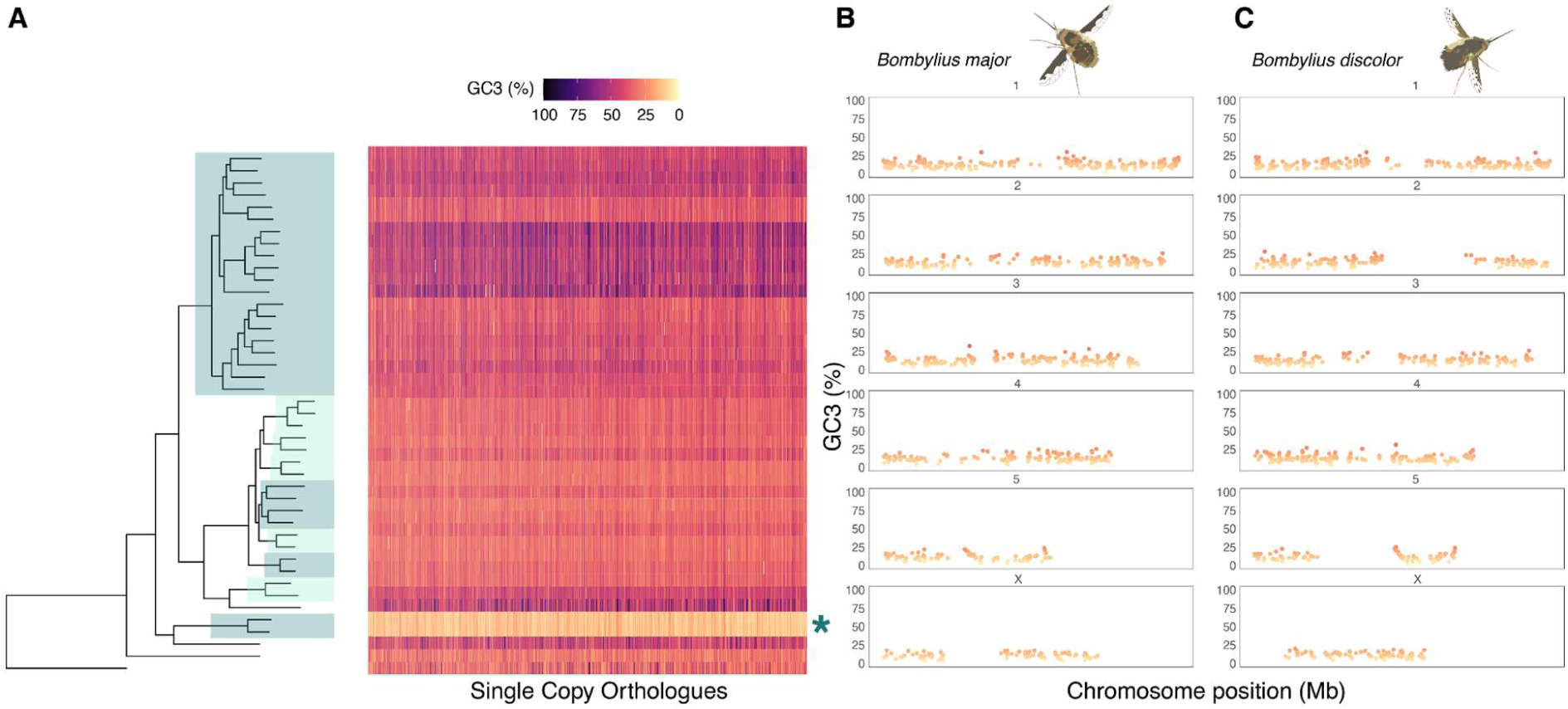
(A) Heatmap showing variation in GC3 for dipteran SCOs across the species tree. SCOs plotted along the x-axis for each species (mapped to species phylogeny) and coloured by GC3 content. Species tree is alternatively highlighted for all families containing two or more representative species. Green asterisk highlights GC3 decrease in *Bombylius* species. (B) Chromosomal landscape plot showing GC3 content (y-axis) of SCOs along chromosomes (x-axis) in *B. major*. (C) Chromosomal landscape plot showing GC3 content (y-axis) of SCOs along chromosomes (x-axis) in *B. discolor*.

**Figure 6.**
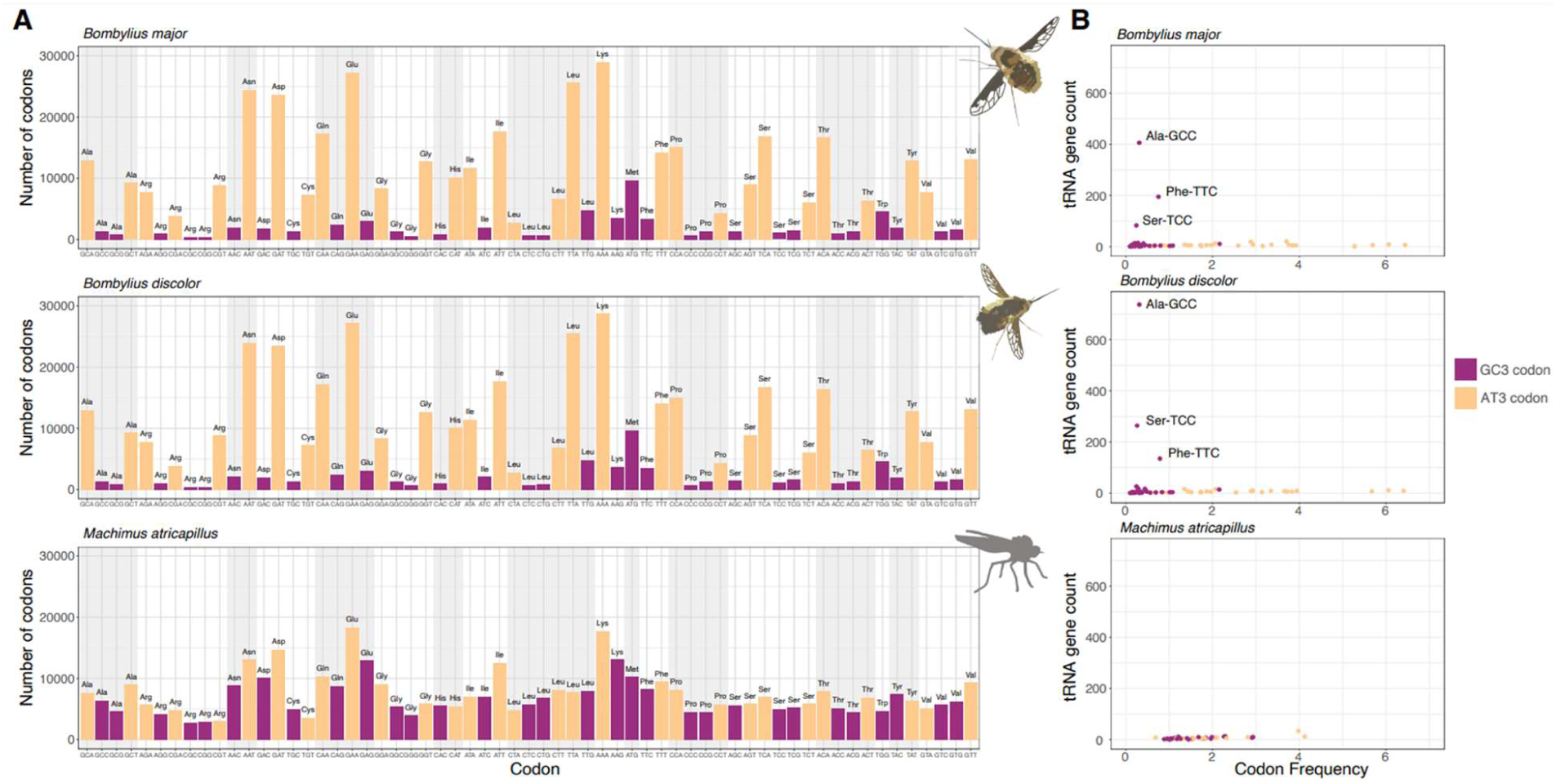
(A) Codon usage (total count, y-axis) in Dipteran SCOs between the *Bombylius major* (top), *Bombylius discolor* (middle) and *Machimus atricapillus* (bottom). In AT3 codons (yellow) the third nucleotide is A or T; in GC3 codons (purple) the third nucleotide is G or C. Stop codons are omitted from the analysis. (B) Plots of tRNA gene counts against codon frequency in Dipteran SCOs. tRNA genes with anticodons binding Ala-GCC, Phe-TTC, and Ser-TCC underwent extreme duplication in *Bombylius* species.

To test if AT-bias was driven by loss of the Base Excision Repair system targeted to deaminated or oxidised cytosine residues, we searched for homologues of *tdg*, *smug1* and *ogg1* genes (Krokan and Bjørås, 2013); homologues of all genes were found in the two *Bombylius* species (Supplementary Figure S3).

We tested if all codons are still used in the AT-rich *Bombylius* genomes by plotting codon usage of the single copy orthologs (Figure 6A); this data set should be immune from artefacts caused due to misprediction of genes or exons as SCO used were aligned at the protein level and trimmed. For comparison, we plotted codon usage for the orthologous genes in the closest relative in the data set, the kite-tailed robberfly, *Machimus atricapillus* (Figure 6A). As expected, the frequency of AT3-codons to GC3-codons is dramatically skewed in *B. major* and *B. discolor*, when compared to *M. atricapillus*. Interestingly, no GC3-codons have been completely omitted from the *Bombylius* genome, implying every codon can be translated.

In some insects, codon usage is associated with transfer RNA (tRNA) abundance (Behura and Severson, 2011; Näsvall et al., 2023b). In *Machimus atricapillus*, we observed the expected result of significant positive correlation between tRNA gene count and codon frequency (Pearson’s product moment correlation coefficient; r = 0.86, P < 1.033e-07; Figure 6B; Supplementary Figure S4), implying more frequent codons in the genome utilise a greater number of tRNAs. However, in both *Bombylius* species no significant positive correlation between tRNA gene count and codon usage is observed (Supplementary Table S4). In both *Bombylius* species, tRNAs with anticodons binding Ala-GCC, Phe-TTC, and Ser-TCC have undergone extensive duplication within the genome (*B. major*: 406, 195, 83; *B. discolor:* 737, 135, 264 respectively), despite these codons being seldom used. This expansion is so extreme that it leads to *B. major* and *B. discolor* containing almost double the amount of predicted tRNA genes compared to all other Diptera in our dataset (Supplementary Figure S5). Paradoxically, *Bombylius* have seen large duplications in three tRNA genes whose anticodons bind some of the least utilised codons in their genomes.

It has been shown previously that, at least in mammals, an accumulation of extreme GC richness can drive aberrant protein divergence and the fixation of deleterious alleles (Dai et al., 2020; Hargreaves et al., 2017). We therefore tested whether extreme AT richness in the *Bombylius* species has driven protein divergence. We used the Sneath index as the measure of ‘relative protein divergence’, restricting analysis to positions in protein sequence alignments that have the same residue in three nested outgroup species (Sneath, 1966) (Figure 7A); such conservation suggests that these residues represent the ancestral state for Diptera and are likely functionally conserved. The Sneath index for each protein tested was divided by protein length to obtain an adjusted Sneath value, which identifies proteins with high rates of protein divergence compared to related species. Comparison between the two *Bombylius* species gives the expected result for two very closely related species: Sneath values are similar and have a high correlation (r = 0.948, p < 2.2e-16; Figure 7B) indicating low protein divergence. When comparing protein divergence between *Bombylius* and *M. atricapillus* the aim of the test was to examine if AT3-rich SCOs are associated with aberrant protein divergence (Figures 7C,D). If this was the case, this would cause deviation in the plot away from the y = x line and towards the x-axis. We do not detect this pattern (Figures 8C,D) suggesting that, despite extreme levels of genome-wide and third codon position AT bias, protein divergence is not significantly affected in these species.

**Figure 7.**
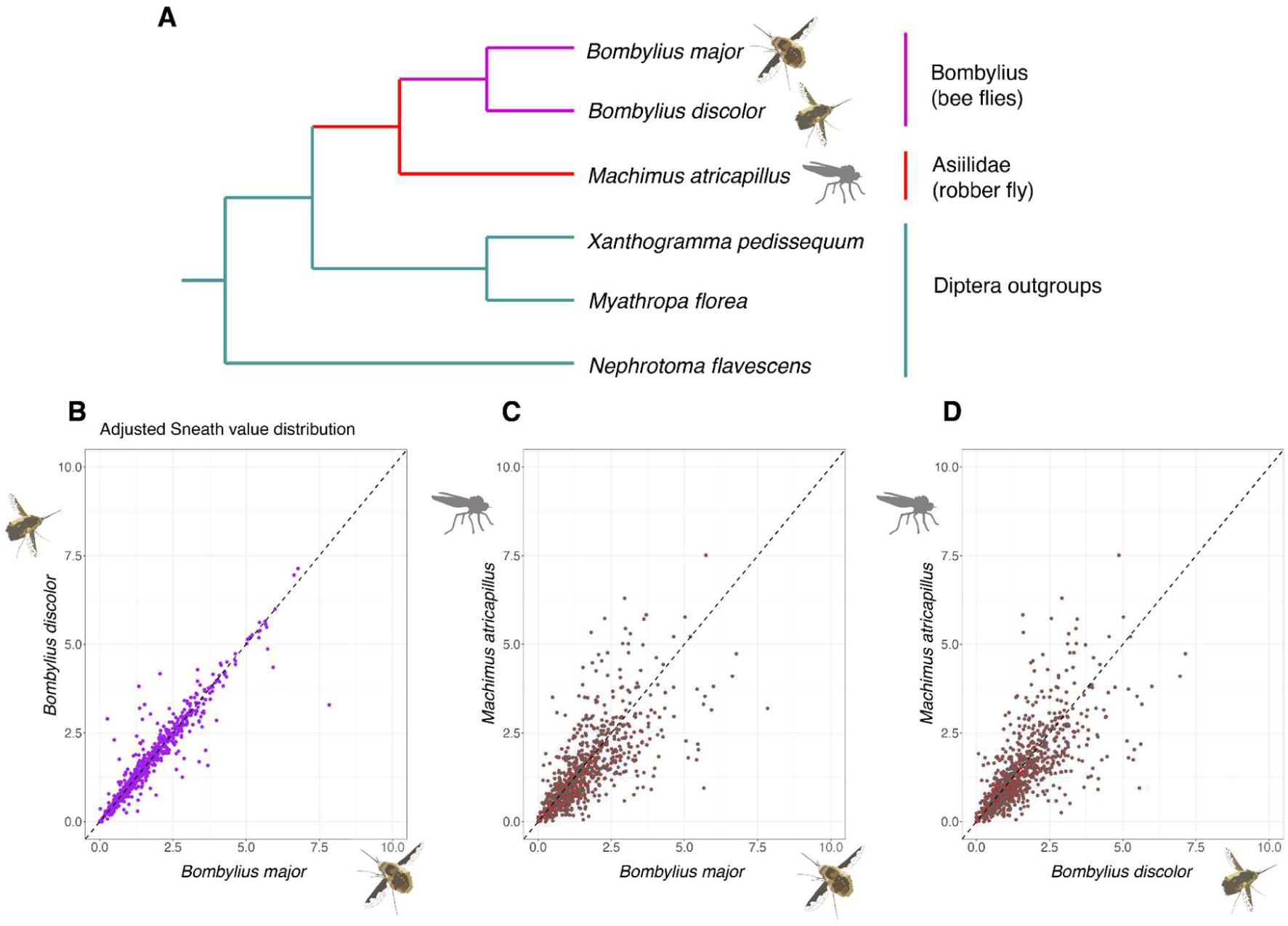
Sneath analysis. (A) Phylogenetic tree showing the two Bombyliidae species and the related *Machimus atricapillus*, each of which is subjected to Sneath analysis based on residues conserved in the three outgroup species. (B) to (D) Plots of adjusted Sneath values compared between species, with each point representing a single SCO. (B) *B. major* vs. *B. discolor.* (C) *M. atricapillus* vs. *B. major.* (D) *M. atricapillu*s vs. *B. discolor*.

## Discussion

### GC-biased gene conversion in holometabolous insects

The incorporation of chromatin capture technology into genome sequencing projects has overcome the challenge of assembling fragmentary de novo genome sequences into chromosomes (Burton et al., 2013; Kaplan and Dekker, 2013; Selvaraj et al., 2013). The 150 insect genomes analysed in the present study have been assembled to chromosomal level; hence, for the first time it is possible to assess GC content in detail along each chromosome in a large number of insect species. A clear trend we uncover is a negative correlation between the percentage GC and distance from the telomere in Lepidoptera, Hymenoptera and Coleoptera (Figure 4). For this analysis, we used GC at the third codon positions of single copy homologous genes, thereby overcoming influences of repetitive DNA or tandemly duplicated genes. We also note that GC content is higher inside protein-coding genes than outside (Figure 1). We suggest that the increase in GC content towards the ends of chromosomes implies that GC-biased gene conversion (gBGC) is prevalent in Lepidoptera, Hymenoptera and Coleoptera genomes. This implication is drawn because gBGC is a recombination-associated event (Duret and Galtier, 2009; Figuet et al., 2014; Pessia et al., 2012; Romiguier and Roux, 2017) and there is evidence that recombination rates increase towards the subtelomeric regions of chromosomes in insects, as in many other taxa (Coop and Przeworski, 2007; Haenel et al., 2018; Ma et al., 2015; Mouresan et al., 2019; Näsvall et al., 2023a; Rockman et al., 2010; Shipilina et al., 2022; Torres et al., 2023).

We note that despite being highly significant, the correlations between chromosome position and GC content have low R squared values, implying data scatter. This does not necessarily imply that gBGC is a minor contributor to the variance in GC3 in insect genomes for three reasons. First, distance from the telomere is an imperfect measure of recombination rate, as each chromosome may have different behaviour in meiosis and may have recombination hotspots at some distance from the telomere. We do not have recombination maps for each species studied. Second, although gBGC is said to be ‘recombination-associated’ this is an over-simplification. In fact, gBGC occurs most frequently when unidirectional strand invasion at meiosis does not lead to a crossover or recombination event, instead resulting in a ‘non-crossover’ or NCO; hence even an accurate recombination map will be an imperfect guide to frequency of gBGC (Duret and Galtier, 2009). Third, we have necessarily used average distance from telomere, and average GC content, for each single copy orthologous gene and this could add some noise to our analyses.

Having argued that gBGC is a likely driver of local GC content in insect genomes, we suggest the strength of this force differs between insect orders. We find that the rate of decline in GC away from the telomere is much steeper in Lepidoptera than in Hymenoptera or Coleoptera, and no decline was detected in Diptera (Figure 4). We suggest, therefore, that gBGC is a stronger force in Lepidoptera. It is not known whether differences in strength of gBGC are caused by biochemical differences, such as the degree of preference for GC over AT by mismatch repair enzymes, or by the length of DNA strands invading at double strand breaks in meiosis, or by frequency of double strand breaks, or other factors. Whatever the proximal reason, the increased strength of gBGC in Lepidoptera is likely to underlie the broad-scale differences in GC content we note between orders, with GC content of coding regions and GC3 of single copy orthologs higher than in other orders (Figure 1).

One prediction of gBGC that is not fulfilled in all species in our analysis is the expectation that smaller chromosomes have higher GC content (Goodstadt et al., 2007; Matsubara et al., 2012). We find this effect in most species of Lepidoptera, but only in a small number of species of Coleoptera, Diptera and Hymenoptera. The effect in Lepidoptera has been noted previously in the butterfly *Vanessa cardui* (Shipilina et al., 2022) and in several other species (Wright et al. 2023). Such an effect is attributed to the greater per-megabase effect gBGC exhibits on smaller chromosomes, particularly in vertebrates, where recombination events encapsulate more of the chromosome proportionally (Goodstadt et al., 2007; Matsubara et al., 2012; Pessia et al., 2012). How can we reconcile evidence for gBGC in Lepidoptera, Hymenoptera and Coleoptera, yet a consistent ‘small chromosome effect’ only in Lepidoptera? We speculate that a possible explanation may lie in numbers of strand invasions per chromosome pair in meiosis. It is recognized that at each meiosis in eukaryotes a minimum of one crossover event must occur between each homologous chromosome pair to facilitate effective segregation. If the number of such events is close to one per chromosome pair, then logically each base pair in a small chromosome will be subject to a strand invasion more frequently than a base pair in a large chromosome. Hence, gBGC would occur more often - per base pair - in smaller chromosomes. In contrast, however, if it is more normal to have multiple strand invasions per chromosome pairing, and these are spaced at regular intervals, then the frequency of gBGC will less strictly dependent on chromosome size. Instead, the rate of gBGC and hence GC content will be dependent on the landscape of double strand breaks and subsequent homologous strand invasion, not only chromosome size.

### Outlier species and the unusual case of bee-flies

We find that GC content of the genome can vary greatly between species within the same order. This intra-order variation is greatest in the Hymenoptera, where we identify three distinct outliers: the Ichneumonidae family of parasitic wasps, the Vespidae family of stinging wasps and the *Bombus* genus of bumblebees. In contrast, Lepidoptera, although having relatively high GC content within coding regions compared to the other insects, has fewer outlier groups or species indicating homogeneity across the order.

The most extreme GC outlier clade in the four insect orders analysed lies in the Diptera and comprises two species of bee-fly in the genus *Bombylius* (family Bombyliidae). The Dark-edged bee-fly *B. major* is a highly active bee mimic found across the northern hemisphere. It is a common spring-flying insect in the UK, often seen hovering to feed on pollen and nectar or to flick eggs into the nests of solitary bees where the bee-fly larvae are predatory. The Dotted bee-fly *B. discolor* is a less common species in the UK, with similar ecology and behaviour to *B. major*. We find that both species have dramatically lower GC content than other Diptera, with the low GC content spanning across all chromosomes and particularly low at third codon positions of protein-coding genes. With a total genome GC content of around 26%, these species are comparable to the most GC-poor genomes found in arthropods, and they represent the most GC-poor dipteran genomes reported to date (Dennis et al., 2020). The low GC content at third codon positions is particularly striking, being below 13% in each species. By comparison, the two most GC-poor arthropod genomes described previously (*Aphidius ervi* and *Lysiphlebus fabarum,* Hymenoptera: Braconidae) have reported GC3 of 15.5 and 10.7%, respectively (Dennis et al., 2020). We stress that the bee-fly figures should not be compared directly to the reported Hymenoptera values as the latter were not based on aligned and trimmed orthologs, so could be more susceptible to annotation inconsistencies.

Three evolutionary questions arise when considering the aberrantly low GC content of bee-fly genomes. When did the reduced GC content evolve? How does it relate to bee-fly biology? How did it evolve? Identifying similar patterns of GC content in two closely related species verifies the legitimacy of the findings and, by considering these species in a phylogenetic context, it is clear that low GC is a derived character for these Diptera. Low GC content of the whole genome, and extremely low GC at third codon positions, therefore evolved on the lineage leading to the genus *Bombylius*, after it had diverged from the lineage leading to the kite-tailed robberfly *Machimus atricapillus.* No additional genome assemblies with the required annotation detail were available to split this evolutionary lineage further, suitable for all the analyses undertaken here. However, one informative genome has been sequenced, although not annotated, and gives some insight: the Downland bee-fly *Villa cingulata,* a more distant member of the Bombyliidae (ENA Assembly GCA_951394055, idVilCing2.1; https://www.ebi.ac.uk/ena/browser/view/GCA_951394055). This has a GC content of 27%, similar to *Bombylius*. We cannot calculate GC3 of single copy orthologs. We therefore suggest that the process that drove the dramatic reduction in GC occurred early in the evolution of the bee-fly lineage.

Relating low GC content to bee-fly biology is difficult. There is no obvious selective reason why the ecology of bee-flies should have been associated with genome-wide reduction of GC content. Other insects also feed on pollen and nectar, have parasitic larvae, or have highly active flight behaviour, and these do not show parallel evolutionary changes. Nitrogen limitation has been proposed as an environmental factor that can cause some selection for lower GC (Acquisti et al., 2009; Foerstner et al., 2005; Šmarda et al., 2014), but this seems insufficient as an explanation for such an extreme, genome-wide case. Currently, therefore, we do not suggest the reduction in GC is adaptive, although the genomic changes may have secondary consequences for bee-fly biology. One consequence could be related to codon usage. Codon bias within *B. discolor* and *B. major* is extremely AT-biased; for every amino acid with redundant codons, the AT3 variant is always utilised in greater numbers. Such extreme AT-codon bias in *Bombylius* is similar to that observed in the AT-rich Hymenoptera *A. ervi* and *L. fabarum* (Dennis et al., 2020), and very different from *Drosophila* and other dipterans (Behura and Severson, 2011; Moriyama and Powell, 1997; Vicario et al., 2007). It is notable that in all four highly AT-rich insect species, all possible codons are utilised within protein-coding genes, suggesting that all codons can be translated, even when not strictly necessary. Codon usage bias can affect translation efficiency and may have negative functional consequences for the species in question (Behura and Severson, 2011; Dennis et al., 2020). Paradoxically, we do not find coevolution between tRNA gene copy number and codon usage, whereby expansion of tRNA genes could aid efficient translation of common codons (Higgs and Ran, 2008; Näsvall et al., 2023). Instead, we find duplication of the genes encoding tRNAs necessary for translating some of the most seldom used codons in *Bombylius* genes.

A second way in which extreme nucleotide bias could affect the biology of bee-flies is if codon positions 1 and 2 are also affected, leading to unusual amino acid substitutions in some proteins. Such a situation has been described in gerbils, where extremely high GC-bias in localised genomic regions has caused fixation of slightly deleterious alleles, recognised through the evolution of unusually divergent proteins (Dai et al., 2020). We have tested for a similar effect in *Bombylius* proteins using a Sneath analysis, and find no evidence that the drive to low GC-content has caused shifts in protein sequence evolution (Figure 7). This is not a perfect test, as the Sneath analysis focussed on single copy orthologous genes, likely to have core insect functions and possibly highly constrained (Conant and Wolfe, 2008). In summary, we suggest the reduction in GC content was not driven by selection, but nor do we find evidence for it adversely affecting protein sequence evolution. The drive to low GC had consequences for genome evolution and may have slightly deleterious consequences through effects on efficiency of mRNA translation. Small effective population size could play a role in permitting fixation of slightly deleterious changes (Galtier et al., 2018).

Finally, we must consider molecular or biochemical mechanisms that might underpin a genome-wide drive towards reduced GC in bee-fly genomes. In many eukaryotes, C:G to T:A mutations can occur either through misincorporation of nucleotides during DNA synthesis, or by deamination of C or methyl-C leading to U:G or T:G pairs which lead to a transition mutation if not recognised and corrected by the Base Excision Repair (BER) process (Krokan and Bjørås, 2013). Since deamination occurs at a high rate, especially in single stranded DNA (Krokan and Bjørås, 2013), this leads to a selective pressure to (a) recognize and correct mismatched base pairs, and (b) counteract the gradual shift towards A:T, for example driving counteracting GC-biased mechanisms such as meiotic gBGC. Theoretically, a trend towards increasing genome-wide AT content could be caused by modification to any of these processes. The deep-sea bone-eating polychaete worm *Osedax frankpressi* has an AT-rich genome similar to that of *Bombylius* (GC content 29%; Moggioli et al., 2023). In this case, an underlying cause is proposed to be loss of genes encoding components of the BER pathway, including SMUG1 which recognizes U:G base pairs resulting from C deamination (Krokan and Bjørås, 2013; Moggioli et al., 2023). However, we suggest this may not be the underlying cause in bee-flies, since we found homologues of BER genes missing from *Osedax*, including *smug1*, *tdg* and *ogg1*, in *Bombylius* and all flies in our data set (Supplementary Figure S3). Although methyl-C is generally low in dipteran genomes, associated with loss of genes encoding DNMT1 and DNMT3 (Provataris et al., 2018), deamination also affects unmethylated cytosine so a rate increase in deamination is a possible cause. Another possibility is that the process of biased gene conversion has become less GC-biased in this evolutionary lineage, although we have no direct evidence to support this speculation. It is therefore possible that the AT-richness in *Bombylius* may have arisen through a complex interplay between biochemical factors such as those discussed above, in concert with effective population size and environmental factors.

## Methods

### Measurements of GC content

Insect genomes and annotation data were primarily by the Darwin Tree of Life project (The Darwin Tree of Life Project Consortium, 2022), apart from *Neomicropteryx cornuta* (Li et al. 2021). Genome data was downloaded from NCBI and annotation data was obtained from the Ensembl rapid release site using publicly available scripts (https://github.com/PeterMulhair/DToL_insects). A list of species (60 Lepidoptera, 42 Diptera, 33 Hymenoptera, and 15 Coleoptera) and their genome accession identifiers study are given in the Supplementary Data associated with this manuscript.

Chromosome GC content was calculated using a python script (chrm_GCcontent.py) for chromosomes with a lower end cut-off of 1Mb to remove unplaced scaffolds, and CDS GC content was calculated using a similar script (cds_GCcontent.py) for chromosome CDS regions excluding gene fragments <950 nucleotides. To generate four sets of single copy orthologs (SCOs), protein FASTA files for all species in each order were downloaded from Ensembl rapid release site and OrthoFinder v2.5.4 was used to build gene families (Emms and Kelly, 2019) for each order. A script from OrthoFinder tools (primary_transcript.py) was used to ensure only primary transcripts were used to construct gene families, and SCOs were automatically generated by OrthoFinder for all four orders. OrthoFinder runs resulted in 1,944 SCOs in Hymenoptera, 1,330 in Coleoptera, 849 in Diptera, and 772 in Lepidoptera. Measurement of third codon position GC (GC3) used the script orthofinder_GC3.py (github.com/RiccardoKyriacou/Insect_GC_content) which aligns all SCOs at the protein level using MAFFT v7.467 (Katoh et al., 2002), trims genes to ensure that only conserved, homologous regions are considered, and back translates to nucleotide sequences to count GC3 using trimAL v1.4 (Capella-Gutiérrez et al., 2009). The mean genome GC, CDS GC, and SCO GC3s were calculated from the output and used to create Figure 1 using ggplot2 in R. A two-way ANOVA was conducted on the data (Supplementary Table S1).

### Phylogenetic plots of insect genome GC, CDS GC, and single copy ortholog GC3 content

Species tree outputs from OrthoFinder runs were used to generate Figure 2; these default trees are based on all orthogroups using the STAG method (Emms and Kelly, 2019). The R packages ggTree and ggTreeExtra were used to generate plots with concentric rings of GC content mapped to the species tree (Xu et al., 2022, 2021).

### Analysis of chromosomal GC content against chromosome size

Using the output from chrm_GCcontent.py that contained chromosome length and GC content for each species, Figure 3 was generated using ggplot2. Only whole chromosomes were considered for this analysis; any contig not mapped to a chromosome was ignored. Regressions were fitted to the data for each order and for each species using a linear model of GC and chromosome size (Supplementary Table S2; Supplementary Data).

### Analysis of GC content and position along the chromosome

By comparing gene ID in the single copy orthologs file to the CDS/genome files, orthofinder_GC3.py retrieves position along the chromosome; script get_telomere_plot.py calculates distance from telomere for each SCO. Genes were plotted individually in Figure 4B and Figure 5B,C; for Figure 4A, we calculated mean GC3 and mean average distance from telomere between all species in an order and conducted regression analysis using R (Supplementary Table S3).

### Assessment of the impact of AT richness in *Bombylius*

The orthofinder_GC3.py script compares GC3 content for each SCO between species in a dataset, and produces an output of orthogroup ID, species, and GC3 values. For Diptera, each orthogroup was plotted on a heatmap coloured by GC3, alongside the species tree, to visualise evolutionary changes in GC3 (Figure 6A).

Calculation of codon usage used script codon_usage.y which, when run on a concatenated file of all SCOs for a species, achieved using concat_OGs.py, calculates an absolute measure of codon usage for that species, and determines whether each codon is AT3 (A/T nucleotide at the third position) or GC3 (G/C nucleotide at the third position); these data were used to plot Figure 6. To calculate adjusted Sneath value (Figure 7B-D) for *Bombylius* SCO proteins, Python script get_sneath_index.py was written to find amino acid residues conserved among the three specified Diptera outgroups. If the residue is conserved, representing the ancestral state, then the equivalent residue is identified for each ingroup species. Adjusted Sneath value is calculated by comparing the ancestral residues to the residues for each ingroup species, according to the Sneath index and dividing the result by the alignment length. Methods for Pearson correlations, Sneath analysis and graphs adapted from Dai et al. (2020) and performed in R (Supplementary Table S5).

To test if any tRNA genes are missing *B. major* and *B. discolor* genome we used tRNAscan-SE v2.0.12 (Chan et al., 2021) with default settings on all dipteran genomes; tRNA-gene models annotated as pseudogenes were excluded from the analyses. BLASTP (Altschul et al., 1990) was used to test if genes encoding TDG, SMUG1 and OGG base excision repair (BER) enzymes are present in *B. major* and *B. discolor* genomes; gene trees were built using IQTREE (Minh et al., 2020).

## Acknowledgements

We thank Lindsay Turnbull, Steve Kelly, Asia Hoile and Liam Crowley for helpful discussions, and all members of the Darwin Tree of Life Consortium for their dedication to making genome sequences freely available to all. This research was funded by the Wellcome Trust Darwin Tree of Life Awards [218328, 226458].

## Data availability

All scripts used during this study can be found in the following GitHub repository, (https://github.com/RiccardoKyriacou/Insect_GC_content) with comments providing code explanations. All data extracted from genome sequences are compiled in the Supplementary Data table associated with this manuscript (genome IDs; genome GC; CDS GC; SCO GC3; species tree files; chromosome size GC; regressions between chromosome size and GC; SCO gene identification numbers, chromosomal location and GC3; average distances from telomeres; codon usage data; Sneath values).

