## Supplementary Materials for "GC content across insect genomes: phylogenetic patterns, causes and consequences"

#### Supplementary Figures

Supplementary Figure S1: Labelled species trees for Lepidoptera, Diptera, Hymenoptera and Coleoptera
Supplementary Figure S2: Chromosome sizes for Lepidoptera, Diptera, Hymenoptera and Coleoptera
Supplementary Figure S3: Amino acid trees for genes encoding OGG1, TDG, SMUG1
Supplementary Figure S4: Codon usage against tRNA correlations, outliers removed
Supplementary Figure S5: tRNA gene count and anticodon GC% across Diptera

#### Supplementary Tables

Supplementary Table S1: Two-way ANOVA and TukeyHSD post-hoc test on GC content with DNA type and order
Supplementary Table S2: GC content against chromosome size regression analysis
Supplementary Table S3: GC content against distance from telomere regression analysis
Supplementary Table S4: tRNA against codon usage analysis
Supplementary Table S5: Sneath value analysis

### Supplementary Figures

Supplementary Figure S1: Labelled species trees for Lepidoptera, Diptera, Hymenoptera and Coleoptera

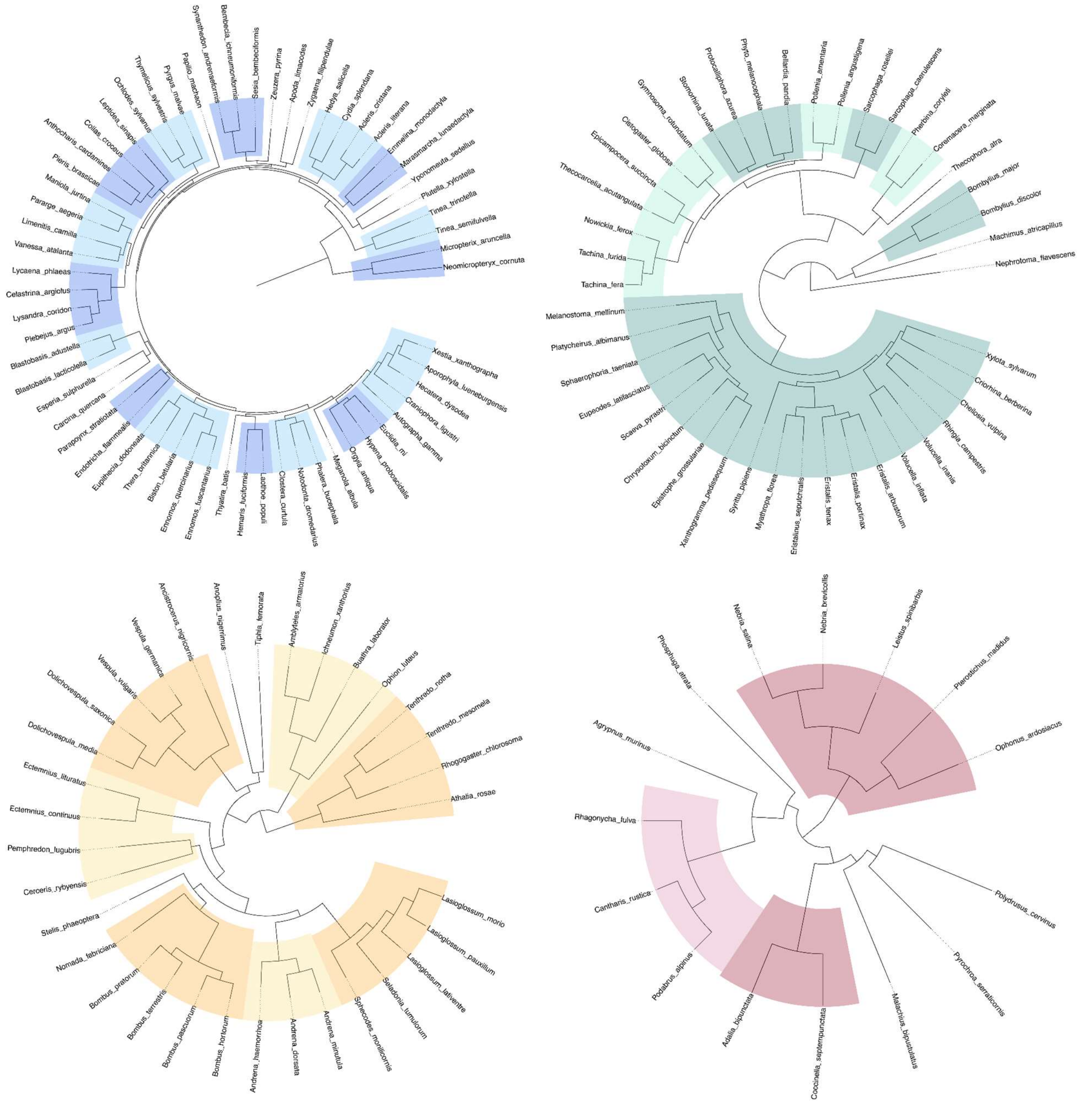

**Supplementary Figure S1. Radial species trees for 150 insects in this study. 60 Lepidoptera (top left), 42 Diptera (top right), 33 Hymenoptera (bottom left), 15 Coleoptera (bottom right). Species trees generated from in OrthoFinder using default settings and STAG methodology.**

Supplementary Figure S2: Chromosome sizes for Lepidoptera, Diptera, Hymenoptera and Coleoptera

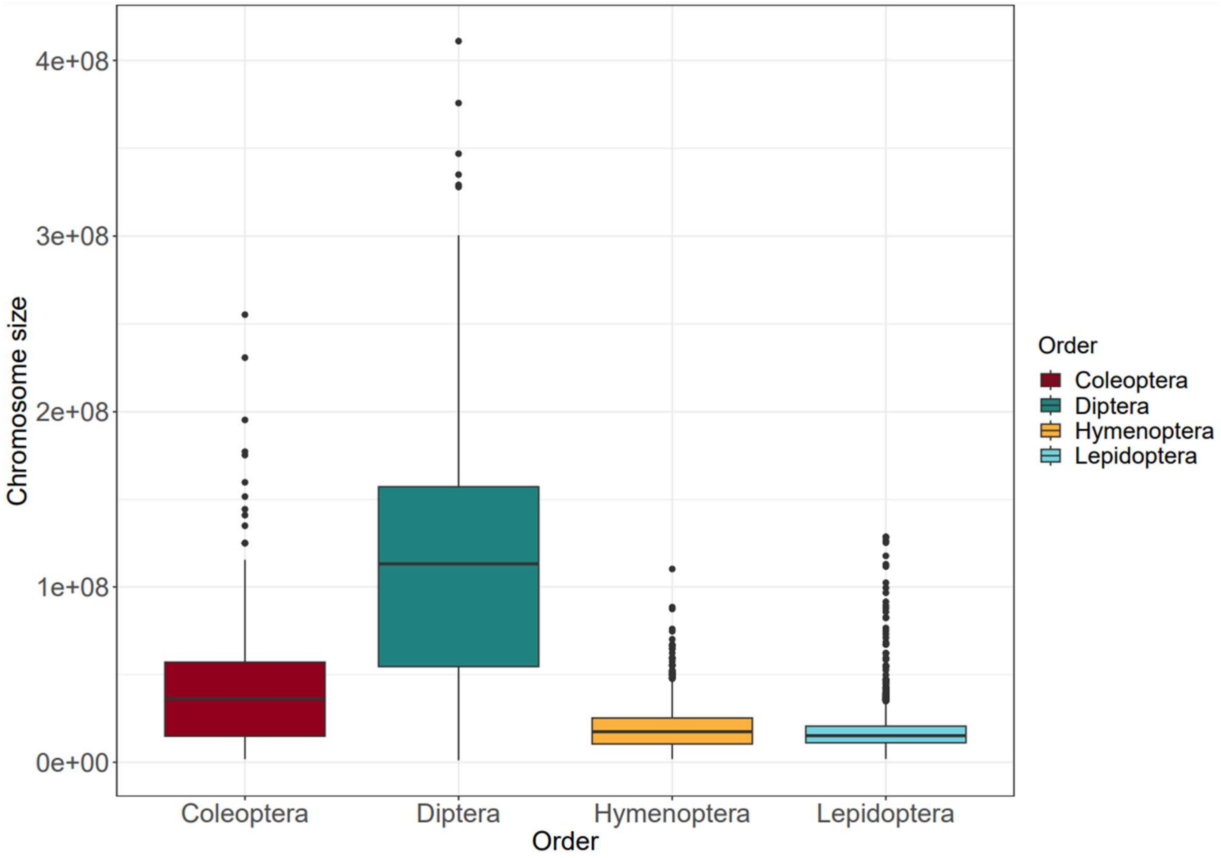


**Supplementary Figure S2. Comparison of chromosome sizes for Coleoptera (red), Diptera (green), Hymenoptera (yellow) and Lepidoptera (light blue) chromosomes. Each chromosome for each species plotted individually for 60 Lepidoptera (1827 chromosomes), 42 Diptera (237 chromosomes), 33 Hymenoptera (446 chromosomes) and 15 Coleoptera (197 chromosomes) species. Box values correspond to the first and third quartiles. Upper whisker extends to the largest value 1.5x interquartile range or distance between the first and third quartiles. Lower whisker extends to the smallest value 1.5 * IQR. Any data points beyond these limits are represented as outlier points.**

### Supplementary Figure S3: Amino acid trees for genes encoding OGG1, TDG, SMUG1

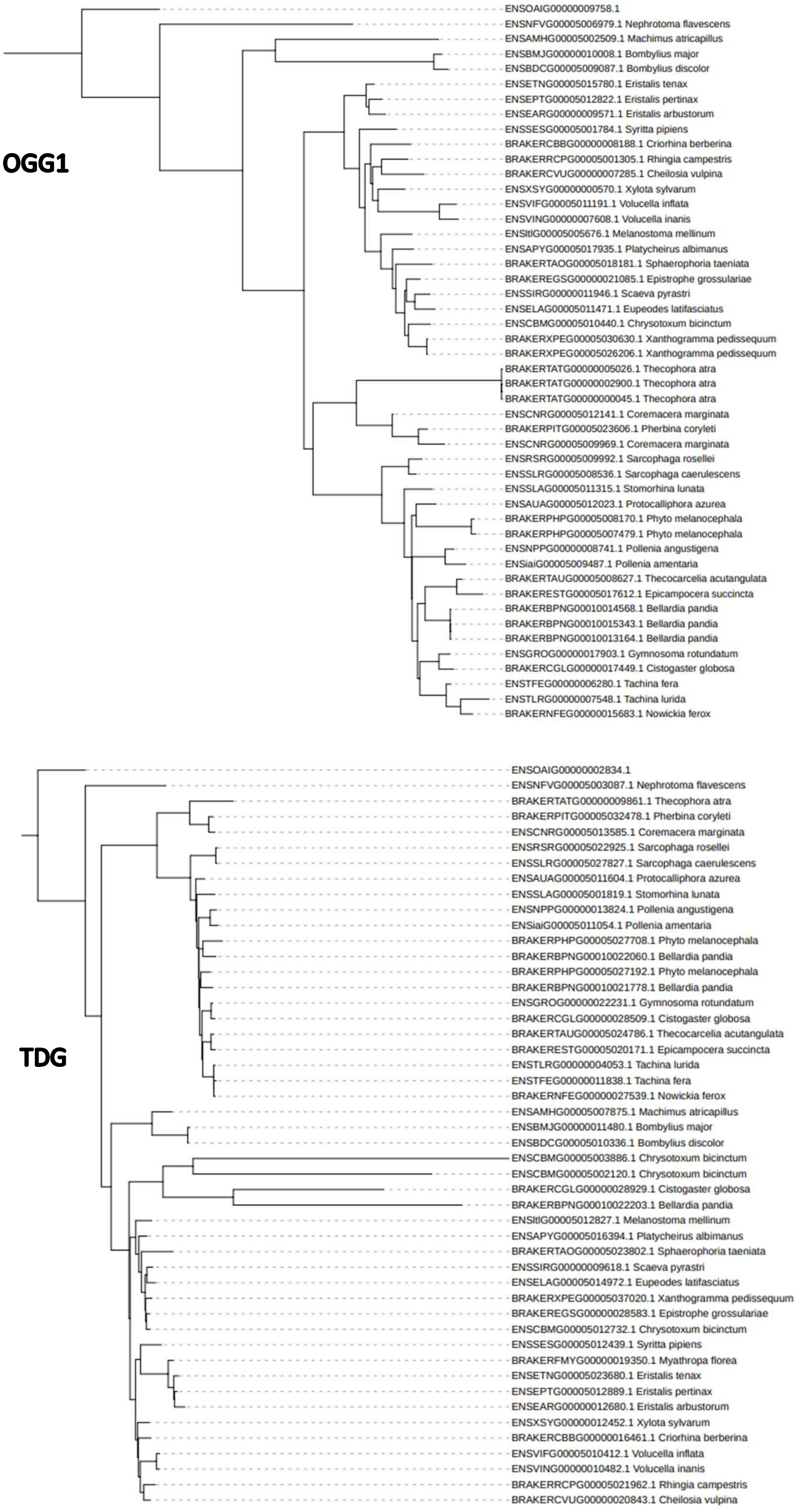

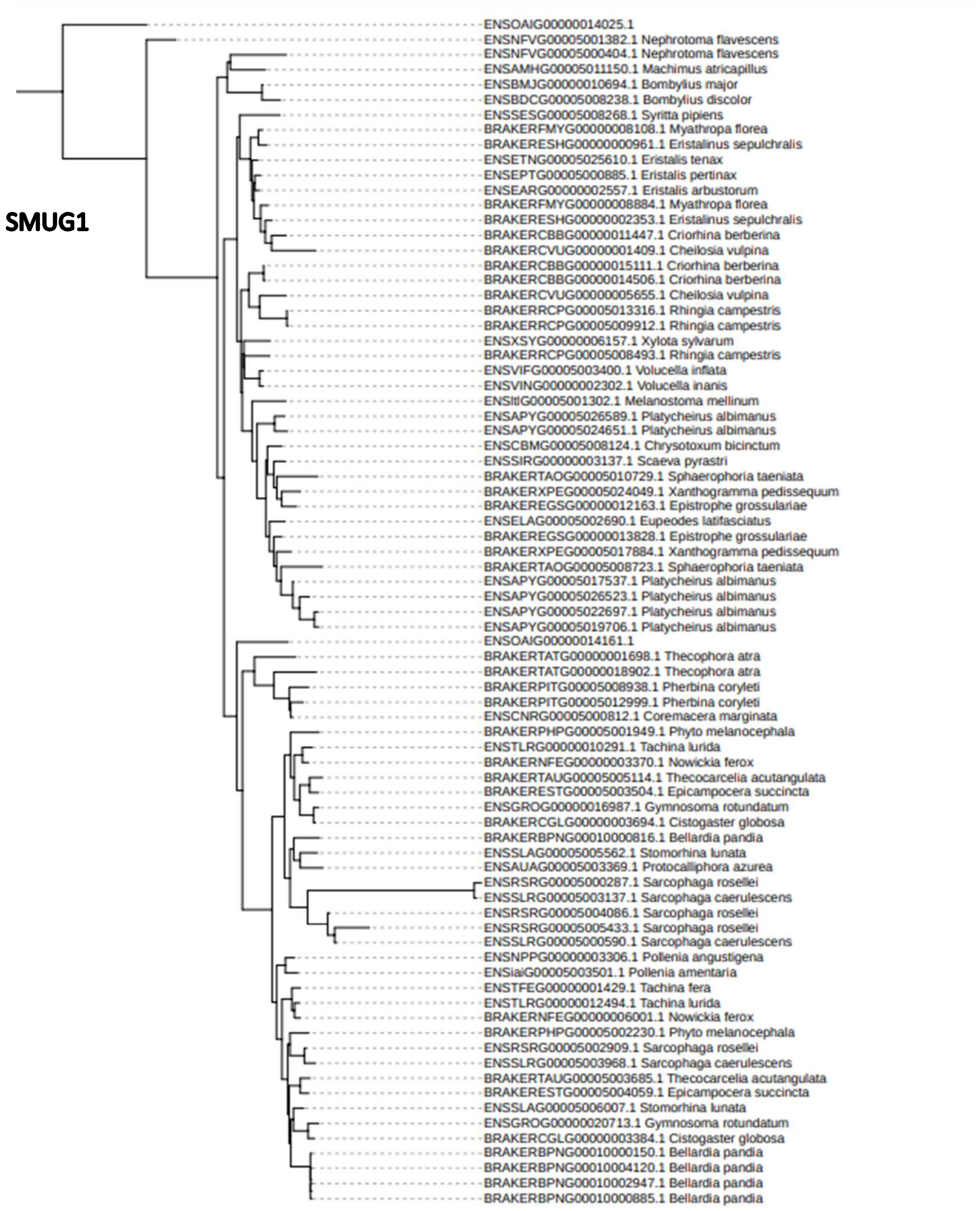


**Supplementary Figure S3 - Gene trees built from amino acid sequences for the Base Excision Repair (BER) genes *ogg1* (top)*, tdg* (middle) and *smug1* (bottom) found across dipteran species (plus one outgroup species, *Orgyia antiqua* Lepidoptera). The OGG1 protein primarily targets oxidizes G resides which can otherwise lead to G:C to A:T mutations; TDG primarily targets T:G pairs resultant from deamination of 5methyl-C residues, but also can target U:G pairs; SMUG1 primarily targets U:G pairs generated by deamination of C residues. Tree tip labels show gene ID followed by species names. Gene trees built using IQTREE.**

Supplementary Figure S4: Codon usage against tRNA correlations with outliers removed

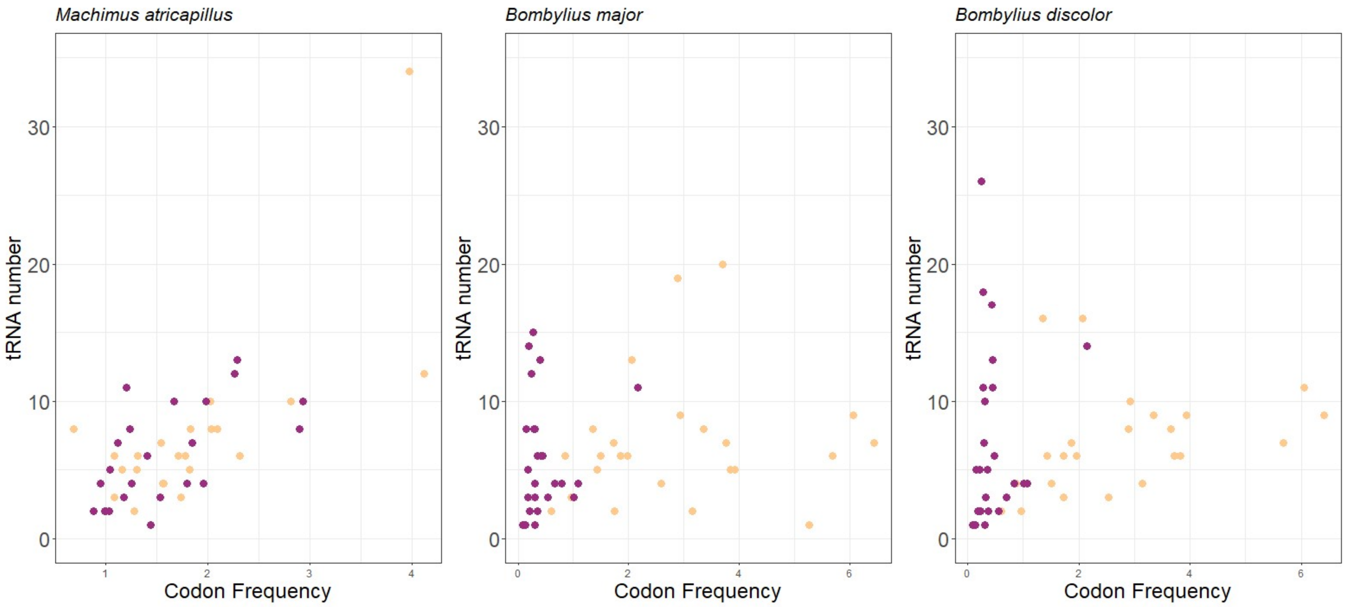


**Supplementary Figure S4 - Plots of tRNA gene counts against codon frequency in Dipteran SCOs. Outliers in *B. major* and *B. discolor* (that is tRNA genes with anticodons binding Ala-GCC, Phe-TTC, and Ser-TCC) have been excluded from the data, in order to better understand the relationship between the two variables.**

Supplementary Figure S5: tRNA gene count and anticodon GC% across Diptera

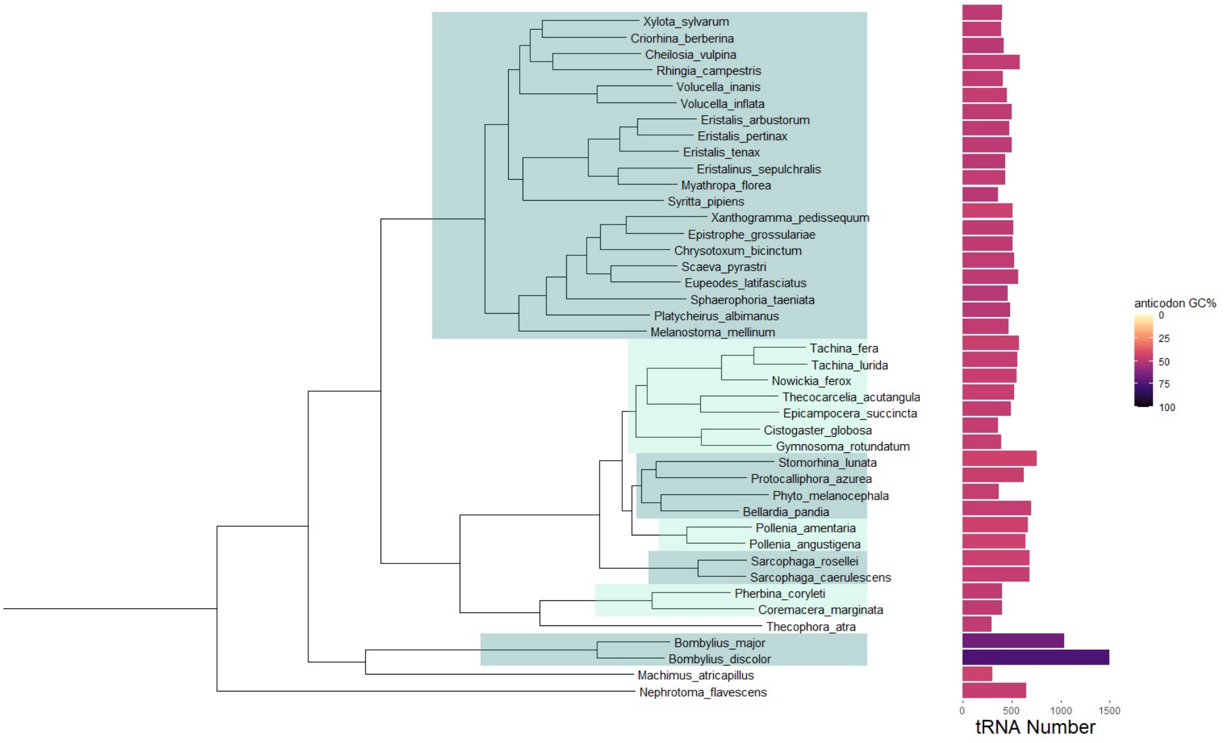


**Supplementary Figure S5 - Dipteran species tree plotted next to bar chart showing tRNA gene number for each species. tRNA numbers predicted from dipteran genomes using tRNAscan-SE 2.0. For each species, bars are coloured according to the average GC content of the tRNA anticodon. *Bombylius major* and *Bombylius discolor* are outliers in both tRNA number and anti-codon GC% (*B. major*: 1033, 66.9%; *B. discolor*: 1493, 75.7%).**

### Supplementary Tables

#### Table S1: Two-way ANOVA and TukeyHSD post-hoc test on GC content with DNA type and order

##### 1a) Two way ANOVA:

m1 <- aov(gc ~ DNA_type * order, data
sp_combined_table)

| | | | | |
|---|---|---|---|---|
| DNA_type | 2 | 2938.4 | 118.26 | < 2e-16 |
| order | 3 | 1632.8 | 65.71 | < 2e-16 |
| DNA_type:order | 6 | 307.8 | 12.39 | 6.66E-13 |
| Residuals | 438 | 10883 | 24.8 | |

Signif. codes:  0 ‘***’ 0.001 ‘**’ 0.01 ‘*’ 0.05 ‘.’ 0.1 ‘ ’ 1

##### 1b) Tukey HSD test

| $DNA_type | | | | |
|---|---|---|---|---|
| | diff | lwr | upr | p adj |
| GC_genome-GC_cds | -8.167 | -9.520604 | -6.8133963 | 0 |
| GC3-GC_cds | -1.126533 | -2.480137 | 0.2270704 | 0.1242083 |
| GC3-GC_genome | 7.040467 | 5.686863 | 8.3940704 | 0 |

| $order | | | | |
|---|---|---|---|---|
| | diff | lwr | upr | p adj |
| Diptera-Coleoptera | 0.9929365 | -1.2395218 | 3.225395 | 0.6605499 |
| Hymenoptera-Coleoptera | 5.6130303 | 3.3018479 | 7.924213 | 0 |
| Lepidoptera-Coleoptera | 8.1088889 | 5.9663655 | 10.251412 | 0 |
| Hymenoptera-Diptera | 4.6200938 | 2.8935992 | 6.346588 | 0 |
| Lepidoptera-Diptera | 7.1159524 | 5.6227592 | 8.609146 | 0 |
| Lepidoptera-Hymenoptera | 2.4958586 | 0.8873437 | 4.104373 | 0.0004282 |

| $`DNA_type:order` | | | | |
|---|---|---|---|---|
| | diff | lwr | upr | p adj |
| GC_genome:Coleoptera-GC_cds:Coleoptera | -7.352667 | -13.333471 | -1.3718626 | 0.0036194 |
| GC3:Coleoptera-GC_cds:Coleoptera | -3.852 | -9.832804 | 2.128804 | 0.6113421 |
| GC_cds:Diptera-GC_cds:Coleoptera | 0.2620952 | -4.6646188 | 5.1888093 | 1 |
| GC_genome:Diptera-GC_cds:Coleoptera | -6.209095 | -11.135809 | -1.2823812 | 0.0024178 |
| GC3:Diptera-GC_cds:Coleoptera | -2.278857 | -7.2055712 | 2.6478569 | 0.9345784 |
| GC_cds:Hymenoptera-GC_cds:Coleoptera | 4.4773333 | -0.6231135 | 9.5777802 | 0.1501339 |
| GC_genome:Hymenoptera-GC_cds:Coleoptera | -0.305697 | -5.4061438 | 4.7947498 | 1 |
| GC3:Hymenoptera-GC_cds:Coleoptera | 1.4627879 | -3.6376589 | 6.5632347 | 0.9986313 |
| GC_cds:Lepidoptera-GC_cds:Coleoptera | 7.6525 | 2.9242593 | 12.380741 | 0.0000108 |
| GC_genome:Lepidoptera-GC_cds:Coleoptera | -3.766333 | -8.4945741 | 0.9619074 | 0.2731306 |
| GC3:Lepidoptera-GC_cds:Coleoptera | 9.2358333 | 4.5075926 | 13.964074 | 0 |
| GC3:Coleoptera-GC_genome:Coleoptera | 3.5006667 | -2.4801374 | 9.4814707 | 0.7441075 |
| GC_cds:Diptera-GC_genome:Coleoptera | 7.6147619 | 2.6880478 | 12.541476 | 0.000036 |

| | | | | |
|---|---|---|---|---|
| GC_genome:Diptera-GC_genome:Coleoptera | 1.1435714 | -3.7831426 | 6.0702855 | 0.9998163 |
| GC3:Diptera-GC_genome:Coleoptera | 5.0738095 | 0.1470955 | 10.000524 | 0.0369518 |
| GC_cds:Hymenoptera-GC_genome:Coleoptera | 11.83 | 6.7295532 | 16.930447 | 0 |
| GC_genome:Hymenoptera-GC_genome:Coleoptera | 7.0469697 | 1.9465229 | 12.147417 | 0.0004479 |
| GC3:Hymenoptera-GC_genome:Coleoptera | 8.8154546 | 3.7150077 | 13.915901 | 0.0000016 |
| GC_cds:Lepidoptera-GC_genome:Coleoptera | 15.005167 | 10.276926 | 19.733407 | 0 |
| GC_genome:Lepidoptera-GC_genome:Coleoptera | 3.5863333 | -1.1419074 | 8.3145741 | 0.347525 |
| GC3:Lepidoptera-GC_genome:Coleoptera | 16.5885 | 11.860259 | 21.316741 | 0 |
| GC_cds:Diptera-GC3:Coleoptera | 4.1140952 | -0.8126188 | 9.0408093 | 0.2086079 |
| GC_genome:Diptera-GC3:Coleoptera | -2.357095 | -7.2838093 | 2.5696188 | 0.9182828 |
| GC3:Diptera-GC3:Coleoptera | 1.5731429 | -3.3535712 | 6.4998569 | 0.9964112 |
| GC_cds:Hymenoptera-GC3:Coleoptera | 8.3293333 | 3.2288865 | 13.42978 | 0.0000085 |
| GC_genome:Hymenoptera-GC3:Coleoptera | 3.546303 | -1.5541438 | 8.6467498 | 0.4891637 |
| GC3:Hymenoptera-GC3:Coleoptera | 5.3147879 | 0.2143411 | 10.415235 | 0.0325504 |
| GC_cds:Lepidoptera-GC3:Coleoptera | 11.5045 | 6.7762593 | 16.232741 | 0 |
| GC_genome:Lepidoptera-GC3:Coleoptera | 0.0856667 | -4.6425741 | 4.8139074 | 1 |
| GC3:Lepidoptera-GC3:Coleoptera | 13.087833 | 8.3595926 | 17.816074 | 0 |
| GC_genome:Diptera-GC_cds:Diptera | -6.47119 | -10.045405 | -2.8969764 | 0.0000004 |
| GC3:Diptera-GC_cds:Diptera | -2.540952 | -6.1151664 | 1.0332617 | 0.452616 |
| GC_cds:Hymenoptera-GC_cds:Diptera | 4.2152381 | 0.4051131 | 8.0253631 | 0.0161071 |
| GC_genome:Hymenoptera-GC_cds:Diptera | -0.567792 | -4.3779172 | 3.2423328 | 0.999998 |
| GC3:Hymenoptera-GC_cds:Diptera | 1.2006926 | -2.6094323 | 5.0108176 | 0.9968036 |
| GC_cds:Lepidoptera-GC_cds:Diptera | 7.3904048 | 4.0951422 | 10.685667 | 0 |

| | | | | |
|---|---|---|---|---|
| GC_genome:Lepidoptera-GC_cds:Diptera | -4.028429 | -7.3236911 | -0.7331661 | 0.0039538 |
| GC3:Lepidoptera-GC_cds:Diptera | 8.9737381 | 5.6784756 | 12.269001 | 0 |
| GC3:Diptera-GC_genome:Diptera | 3.9302381 | 0.3560241 | 7.5044521 | 0.0173798 |
| GC_cds:Hymenoptera-GC_genome:Diptera | 10.686429 | 6.8763036 | 14.496554 | 0 |
| GC_genome:Hymenoptera-GC_genome:Diptera | 5.9033983 | 2.0932733 | 9.7135232 | 0.0000339 |
| GC3:Hymenoptera-GC_genome:Diptera | 7.6718831 | 3.8617582 | 11.482008 | 0 |
| GC_cds:Lepidoptera-GC_genome:Diptera | 13.861595 | 10.566333 | 17.156858 | 0 |
| GC_genome:Lepidoptera-GC_genome:Diptera | 2.4427619 | -0.8525006 | 5.7380244 | 0.3841783 |
| GC3:Lepidoptera-GC_genome:Diptera | 15.444929 | 12.149666 | 18.740191 | 0 |
| GC_cds:Hymenoptera-GC3:Diptera | 6.7561905 | 2.9460655 | 10.566315 | 0.0000007 |
| GC_genome:Hymenoptera-GC3:Diptera | 1.9731602 | -1.8369648 | 5.7832851 | 0.8668566 |
| GC3:Hymenoptera-GC3:Diptera | 3.741645 | -0.06848 | 7.55177 | 0.0596154 |
| GC_cds:Lepidoptera-GC3:Diptera | 9.9313571 | 6.6360946 | 13.22662 | 0 |
| GC_genome:Lepidoptera-GC3:Diptera | -1.487476 | -4.7827387 | 1.8077863 | 0.9445787 |
| GC3:Lepidoptera-GC3:Diptera | 11.51469 | 8.219428 | 14.809953 | 0 |
| GC_genome:Hymenoptera-GC_cds:Hymenoptera | -4.78303 | -8.8152876 | -0.7507731 | 0.0062418 |
| GC3:Hymenoptera-GC_cds:Hymenoptera | -3.014545 | -7.0468027 | 1.0177118 | 0.3705268 |
| GC_cds:Lepidoptera-GC_cds:Hymenoptera | 3.1751667 | -0.3745941 | 6.7249274 | 0.1310285 |
| GC_genome:Lepidoptera-GC_cds:Hymenoptera | -8.243667 | -11.793427 | -4.6939059 | 0 |
| GC3:Lepidoptera-GC_cds:Hymenoptera | 4.7585 | 1.2087393 | 8.3082608 | 0.0008073 |
| GC3:Hymenoptera-GC_genome:Hymenoptera | 1.7684849 | -2.2637724 | 5.8007421 | 0.9546711 |
| GC_cds:Lepidoptera- | 7.958197 | 4.4084362 | 11.507958 | 0 |

| | | | | |
|---|---|---|---|---|
| GC_genome:Hymenoptera | | | | |
| GC_genome:Lepidoptera-GC_genome:Hymenoptera | -3.460636 | -7.0103971 | 0.0891244 | 0.0638427 |
| GC3:Lepidoptera-GC_genome:Hymenoptera | 9.5415303 | 5.9917696 | 13.091291 | 0 |
| GC_cds:Lepidoptera-GC3:Hymenoptera | 6.1897121 | 2.6399514 | 9.7394729 | 0.0000012 |
| GC_genome:Lepidoptera-GC3:Hymenoptera | -5.229121 | -8.778882 | -1.6793605 | 0.0001135 |
| GC3:Lepidoptera-GC3:Hymenoptera | 7.7730455 | 4.2232847 | 11.322806 | 0 |
| GC_genome:Lepidoptera-GC_cds:Lepidoptera | 11.418833 | -14.409235 | -8.4284313 | 0 |
| GC3:Lepidoptera-GC_cds:Lepidoptera | 1.5833333 | -1.4070687 | 4.5737354 | 0.8487369 |
| GC3:Lepidoptera-GC_genome:Lepidoptera | 13.002167 | 10.011765 | 15.992569 | 0 |

**1c) Levene's Test for Homogeneity of Variance and normality checks**

| | Df | F value | Pr(>F) | |
|---|---|---|---|---|
| group | 11 | 11.986 | < 2.2e-16 | *** |
| | 438 | | | |

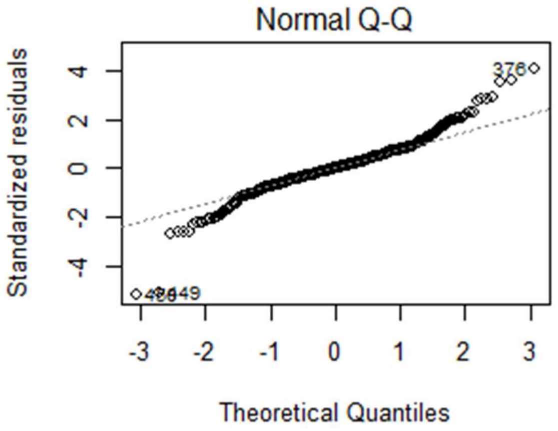


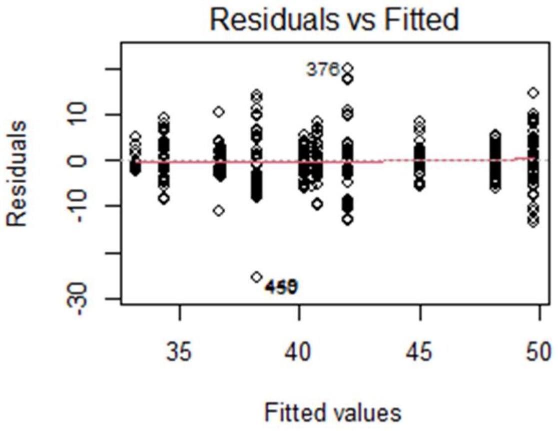

### Table S2: GC content against chromosome size regression analysis

**2a) ANCOVA**
genome_lm <- lm(gc ~ size + order, New_insect_genomes)
Signif. codes: 0 ‘***’ 0.001 ‘**’ 0.01 ‘*’ 0.05 ‘.’ 0.1 ‘ ’ 1

| | Df | Sum Sq | Mean Sq | F value | Pr(>F) |
|---|---|---|---|---|---|
| size | 1 | 1183.1 | 1183.12 | 202.75 | < 2.2e-16 *** |
| order | 3 | 5058.8 | 1686.27 | 288.97 | < 2.2e-16 *** |
| Residuals | 2702 | 15767.2 | 5.84 | | |

**2b) GC vs chromosome size: Lepidoptera**
reg <- lm(gc ~ size, New_insect_genomes%>% filter(order == "Lepidoptera"))

| | | | | |
|---|---|---|---|---|
| Coefficients: | | | | |
| | Estimate | Std. Error | t value | Pr(>\|t\|) |
| (Intercept) | 3.65E+01 | 6.78E-02 | 537.82 | < 2e-16 *** |
| size | 1.83E-08 | 3.17E-09 | 5.77 | 9.27e-09 *** |

Residual standard error: 1.653 on 1825 degrees of freedom
Multiple R-squared: 0.01792, Adjusted R-squared: 0.01738
F-statistic: 33.3 on 1 and 1825 DF, p-value: 9.27e-09, positive slope

**2c) GC vs chromosome size: Coleoptera**
reg <- lm(gc ~ size, New_insect_genomes%>% filter(order == "Coleoptera"))

| | | | | |
|---|---|---|---|---|
| Coefficients: | | | | |
| | Estimate | Std. Error | t value | Pr(>\|t\|) |
| (Intercept) | 3.31E+01 | 2.13E-01 | 155.51 | <2e-16 *** |
| size | 2.34E-09 | 3.44E-09 | 0.68 | 0.497 |

Residual standard error: 2.029 on 195 degrees of freedom
Multiple R-squared: 0.002367, Adjusted R-squared: -0.002749
F-statistic: 0.4626 on 1 and 195 DF, p-value: 0.4972, not significant

**2d) GC vs chromosome size: Diptera**

reg <- lm(gc  ~ size, New_insect_genomes%>% filter(order == "Diptera"))

| Coefficients: | | | | |
|---|---|---|---|---|
| | Estimate | Std. Error | t value | Pr(>\|t\|) |
| (Intercept) | 3.55E+01 | 5.07E-01 | 69.999 | < 2e-16 *** |
| size | -9.67E-09 | 3.61E-09 | -2.677 | 0.00795 ** |

Residual standard error: 4.499 on 235 degrees of freedom
Multiple R-squared:  0.02959, Adjusted R-squared:  0.02546
F-statistic: 7.167 on 1 and 235 DF, p-value: 0.007951, negative slope

**2e) GC vs chromosome size: Hymenoptera**

reg <- lm(gc  ~ size, New_insect_genomes%>% filter(order == "Hymenoptera"))

| Coefficients: | | | | |
|---|---|---|---|---|
| | Estimate | Std. Error | t value | Pr(>\|t\|) |
| (Intercept) | 3.76E+01 | 2.53E-01 | 148.475 | < 2e-16 *** |
| size | 7.37E-08 | 9.90E-09 | 7.438 | 5.31e-13 *** |

Residual standard error: 3.159 on 444 degrees of freedom
Multiple R-squared:  0.1108, Adjusted R-squared:  0.1088
F-statistic: 55.33 on 1 and 444 DF, p-value: 5.31e-13, positive slope

### Table S3: GC content against distance from telomere regression analysis

**3a) Lepidoptera**
regression <- lm(GC3 ~ as.numeric(t_distance), data = mean_distance)
Signif. codes:  0 ‘***’ 0.001 ‘**’ 0.01 ‘*’ 0.05 ‘.’ 0.1 ‘ ’ 1

| Coefficients: | | | | |
|---|---|---|---|---|
| | Estimate | Std. Error | t value | Pr(>\|t\|) |
| (Intercept) | 6.00E+01 | 1.07E+00 | 55.83 | <2e-16 *** |
| (t_distance) | -1.87E-06 | 1.80E-07 | -10.35 | <2e-16 *** |

Residual standard error: 11.11 on 770 degrees of freedom
Multiple R-squared:  0.1222
Adjusted R-squared:  0.1211
F-statistic: 107.2 on 1 and 770 DF,  p-value: < 2.2e-16

**3b) Coleoptera**
regression <- lm(GC3 ~ as.numeric(t_distance), data = mean_distance)

| Coefficients: | | | | |
|---|---|---|---|---|
| | Estimate | Std. Error | t value | Pr(>\|t\|) |
| (Intercept) | 4.20E+01 | 4.56E-01 | 91.987 | < 2e-16 *** |
| (t_distance) | -2.64E-07 | 3.28E-08 | -8.058 | 1.71e-15 *** |

Residual standard error: 5.478 on 1328 degrees of freedom
Multiple R-squared:  0.04662
Adjusted R-squared:  0.0459
F-statistic: 64.94 on 1 and 1328 DF,  p-value: 1.706e-15

**3c) Hymenoptera**
regression <- lm(GC3 ~ as.numeric(t_distance), data = mean_distance)

| Coefficients: | | | | |
|---|---|---|---|---|
| | Estimate | Std. Error | t value | Pr(>\|t\|) |
| (Intercept) | 4.91E+01 | 1.19E+00 | 41.327 | < 2e-16 *** |
| (t_distance) | -9.63E-07 | 1.77E-07 | -5.431 | 6.32e-08 *** |

Residual standard error: 8.647 on 1942 degrees of freedom
Multiple R-squared: 0.01496
Adjusted R-squared: 0.01445
F-statistic: 29.49 on 1 and 1942 DF, p-value: 6.321e-08

**3d) Diptera**

| Coefficients: | | | | |
|---|---|---|---|---|
| | Estimate | Std. Error | t value | Pr(>\|t\|) |
| (Intercept) | 3.88E+01 | 4.40E-01 | 88.091 | <2e-16 *** |
| (t_distance) | -2.20E-08 | 1.42E-08 | -1.548 | 0.122 |

Residual standard error: 4.691 on 847 degrees of freedom
Multiple R-squared: 0.002823
Adjusted R-squared: 0.001645
F-statistic: 2.398 on 1 and 847 DF, p-value: 0.1219

### Table S4: tRNA against codon usage analysis

3a) *Bombylius major*

Pearson's product-moment correlation
data:  trna_codon$tRNA_count and trna_codon$abs_freq
**t = -0.94885, df = 50, p-value = 0.3473**
alternative hypothesis: true correlation is not equal to 0
95 percent confidence interval:
**-0.3916799  0.1451740**
sample estimates:
cor
**-0.1329954**

3b) *Bombylius discolor*

Pearson's product-moment correlation
data:  trna_codon$tRNA_count and trna_codon$abs_freq
**t = -0.98025, df = 50, p-value = 0.3317**
alternative hypothesis: true correlation is not equal to 0
95 percent confidence interval:
**-0.3953982  0.1408642**
sample estimates:
cor
**-0.1373147**

3c) *Machimus atricapillus*

Pearson's product-moment correlation
data:  trna_codon$tRNA_count and trna_codon$abs_freq
**t = -0.98025, df = 50, p-value = 0.3317**
alternative hypothesis: true correlation is not equal to 0
95 percent confidence interval:
**-0.3953982  0.1408642**
sample estimates:
cor
**-0.1373147**

### Table S5: Sneath value analysis

5a) Checking normality

Shapiro-Wilk normality test:

data:  sneath_data$Bombylius_major
**W = 0.884, p-value < 2.2e-16**
data:  sneath_data$Bombylius_discolor
**W = 0.9011, p-value < 2.2e-16**
data:  sneath_data$Machimus_atricapillus
**W = 0.87683, p-value < 2.2e-16**

5b) *Bombylius majo*r vs *Bombylius discolor*

Pearson's product-moment correlation

data:  sneath_data$Bombylius_major and sneath_data$Bombylius_discolor
**t = 86.776, df = 847, p-value < 2.2e-16**
alternative hypothesis: true correlation is not equal to 0
95 percent confidence interval:
 **0.9408323 0.9544937**
sample estimates:
      cor
**0.948098**

5b) *Bombylius majo*r vs *Machimus atricapillus*

Pearson's product-moment correlation

data:  sneath_data$Bombylius_major and sneath_data$Machimus_atricapillus
**t = 29.102, df = 847, p-value < 2.2e-16**
alternative hypothesis: true correlation is not equal to 0
95 percent confidence interval:
 **0.6717655 0.7392046**
sample estimates:
      cor
**0.7070893**

5b) *Bombylius discolor* vs *Machimus atricapillus*

Pearson's product-moment correlation

data:  sneath_data$Bombylius_discolor and sneath_data$Machimus_atricapillus
**t = 29.08, df = 847, p-value < 2.2e-16**
alternative hypothesis: true correlation is not equal to 0
95 percent confidence interval:
 **0.6714693 0.7389597**
sample estimates:
      cor
**0.7068193**
